# Multiplexed dynamic control of temperature to probe and observe mammalian cells

**DOI:** 10.1101/2024.02.18.580877

**Authors:** William Benman, Pavan Iyengar, Thomas Mumford, Zikang Huang, Lukasz J. Bugaj

## Abstract

Temperature is aa critical parameter for biological function, yet there is a lack of approaches to modulate the temperature of biological specimens in a dynamic and high-throughput manner. We present the thermoPlate, a device for programmable control of temperature in each well of a 96-well plate, in a manner compatible with mammalian cell culture and live cell imaging. The thermoPlate maintains precise feedback control of temperature patterns independently in each well, with minutes-scale heating and cooling through ΔT ∼15-20°C. A computational model that predicts thermal diffusion guides optimal design of heating protocols. The thermoPlate allowed systematic characterization of both synthetic and natural thermo-responsive systems. We first used the thermoPlate in conjunction with live-cell microscopy to characterize the rapid temperature-dependent phase separation of a synthetic elastin-like polypeptide (ELP53). We then measured stress granule (SG) formation in response to heat stress, observing differences in SG dynamics with each increasing degree of stress. We observed adaptive formation of SGs, whereby SGs formed but then dissolved in response to persistent heat stress (> 42°C). SG adaptation revealed a biochemical memory of stress that depended on both the time and temperature of heat shock. Stress memories continued to form even after the removal of heat and persisted for 6-9 hours before dissipating. The capabilities and open-source nature of the thermoPlate will empower the study and engineering of a wide range of thermoresponsive phenomena.

## Introduction

Temperature plays important roles in the function and control of biological systems. In humans, core temperature is narrowly maintained around 37°C and, although temperature can rise during infection, the febrile response is also tightly controlled to remain below ∼41°C^1^. In the biosciences, temperature can be harnessed as a non-invasive perturbation, for example with temperature-sensitive protein mutants that provide conditional knockdown of proteins^2–4^.

Separately, heat-shock promoters can allow expression of custom transgenes in response to a brief increase in temperature. Such technologies can control gene expression in model organisms^5^ and, because heat effectively penetrates opaque tissues, also provide remote control of engineered cells within mammals^6–8^. Exposing mammalian cells to temperatures above ∼42°C induces heat shock, which can be toxic over extended periods. This response has been harnessed for cancer therapy, where thermal ablation of tumors is a common, minimally-invasive treatment strategy^9^.

Despite its importance, there are relatively few experimental methods to systematically manipulate temperature. Incubators can be set to desired temperatures, but changes in temperatures are slow, challenging studies of phenomena that occur over fast (∼minutes) timescales. Incubators are also expensive, have a large footprint, and allow only one temperature to be tested at a time. Commercial devices such as the CherryTemp or VAHEAT are similarly limited in throughput. Thermocyclers can be used to specify multiple temperatures simultaneously^10^, but these are not compatible with long term cell survival or simultaneous imaging. The current lack of methods for microscope-compatible temperature modulation in microwell plates limits our understanding of biological responses to temperature changes and hinders the engineering of thermally-controllable biological systems.

We address this gap with the thermoPlate, a device for programmable heating and thermometry of samples in 96-well plates. The thermoPlate allows heating of individual wells with arbitrary temperature profiles and is compatible with both live-cell microscopy and standard cell culture incubators. We characterize thermoPlate performance, finding accurate heating across the plate to < 0.1°C of the desired set point. We also present a model that predicts heat diffusion across the sample plate, allowing the user to rapidly test the feasibility of arbitrary desired heating programs *in silico*. We then harnessed the utility of the thermoPlate to characterize the rapid temperature-dependent phase-separation of an elastin-like polypeptide (ELP). Finally, we examined the dynamics of stress granule (SG) formation in response to graded levels of heat stress. We found that SGs form but spontaneously dissolve due to the formation of biochemical memories of stress, which depended on both the timing and intensity of heat stress.

## Results

### thermoPlate Design, Calibration, and Dynamics

The thermoPlate maintains a desired temperature by continuous deposition and measurement of heat in each well of a 96-well plate (**Figure 1A**). This is achieved with a custom-designed circuit board that accommodates a pair of thermistors in each of 96 positions in the format of a 96-well plate. A thermistor is a resistor whose resistance changes as a function of temperature, thus enabling its common use as a thermometer (“reader”). However, thermistors can also deposit heat when sufficient current is passed through them (“heater”). By pairing one reader and one heater thermistor in each well, the thermoPlate achieves independent measurement and heating of all 96 wells (**Figure 1A, B**). To operate, the user uploads the desired heating protocols to the on-board Arduino microcontroller. The Arduino regulates the current of each heater through one of 12 shift registers linked to 96 transistors, and it receives the temperature readings from each reader connected to a voltage divider through one of 6 multiplexers (**Figure S1**). With the set point and temperature of each well, the Arduino implements proportional-integral-derivative (PID) feedback control, adjusting the current of each heater to reach and maintain the sample at the desired set point for every well (**Figure 1C**. See **Methods** for more details on feedback control). The thermoPlate sits atop a 96-well culture plate to which it is mated with a 3D printed adapter (**Figure 1D**). The thermoPlate can operate in standard cell culture incubators. Alternatively. when the culture plate has a transparent bottom, this assembly is also compatible with live cell imaging using an inverted microscope. The thermoPlate can run in a standalone format but can also be connected to a computer to view real-time readouts of the current well temperatures.

**Figure 1.**
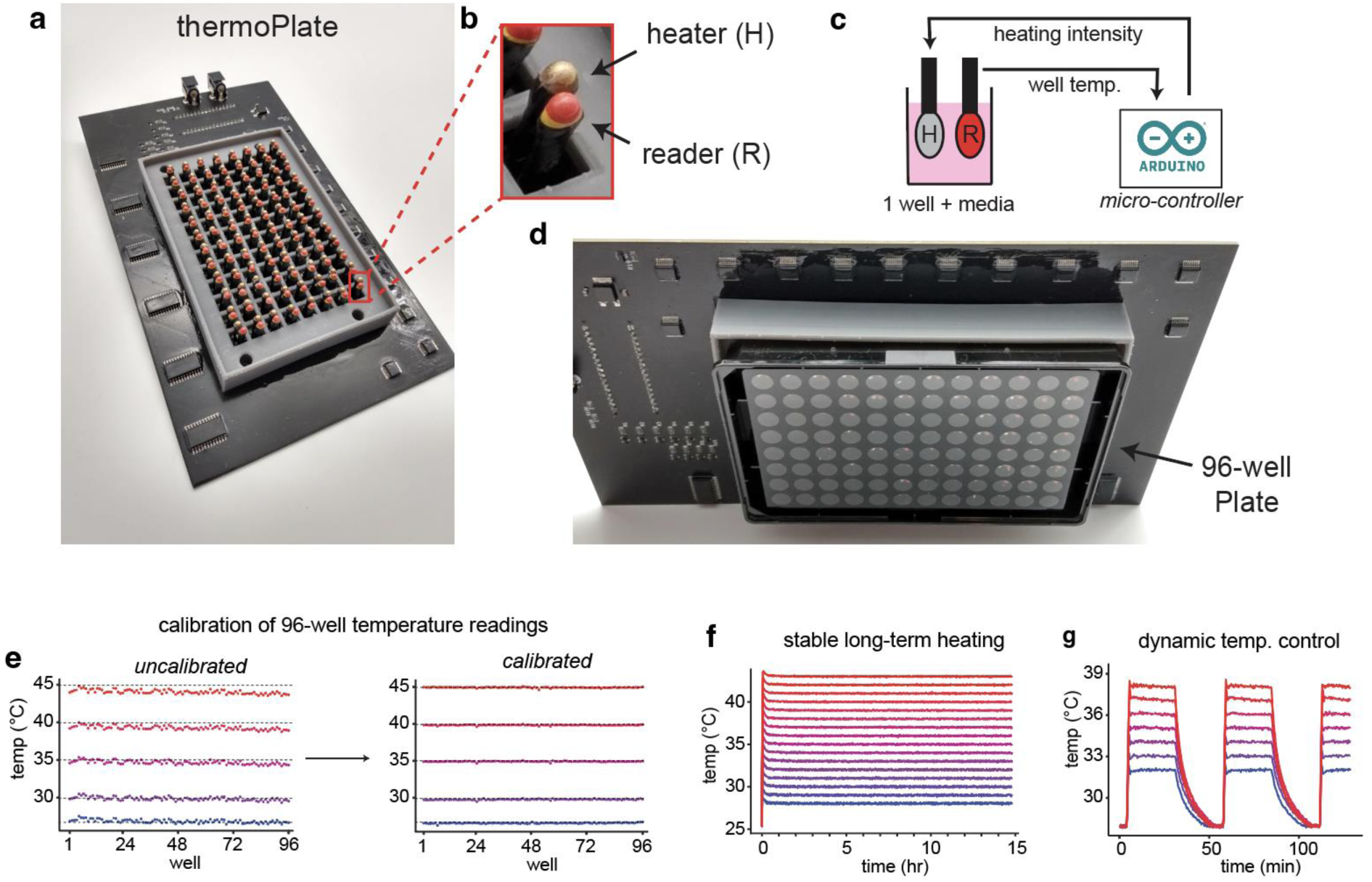
Design and use of the thermoPlate for simultaneous, independent temperature control in 96 well plates. a) Image of a fully assembled thermoPlate device. b) Enlarged image of a single heater/reader pair, which regulates the temperature of a single well of a 96 well plate. c) An Arduino takes readings from a reader and dynamically adjusts the duty ratio of the heater using PID feedback control. e) Raw readings taken from the reader of each well when placed in a cell culture incubator set to various temperatures before and after calibration. Color and dashed lines represent the incubator temperature. See Methods for details on calibration. f) Traces showing 16 wells that were set to 16 different temperatures in 1°C increments (28-43°C) with an ambient temperature of 25°C for 15 hrs. g) Dynamic temperature control in multiple wells over multiple temperatures. 7 wells were heated to different temperatures for repeated cycles of 30 minutes of heating followed by 10 minutes of cooling to 28°C, with an ambient temperature of 25°C.

With all components and 3D printed parts in hand, the thermoPlate can be assembled in 8 hours, for a total cost of ∼$300. Design files, a parts list, assembly instructions, and operation protocols can be found in the **Methods** section and our online repository (link in **Methods**).

Multiplexed temperature control requires consistent, calibrated readings between each of the 96 readers. We characterized the variability in temperature readings in a fully-assembled thermoPlate in a cell culture incubator that was set to one of 5 temperatures between 26-45°C, with the true temperature verified by two digital thermometers. While the average error between thermoPlate and true temperature increased slightly at higher temperatures, we observed a consistent pattern of variability between wells (SD ∼ 0.26°C) (**Figure 1E**). From these results, we derived adjustment factors for each well that corrected readings to within 0.1°C of the true temperature across all temperatures, with a standard deviation of 0.06°C between wells in subsequent experiments (**Figure 1E**). Once calculated, calibration values maintained accurate readings when tested after weeks of use (**Figure S2**).

We next tested the thermoPlate’s ability to control prespecified temperatures, with temperature measured by the calibrated reader thermistors. 15 wells were set to simultaneously maintain temperatures ranging from 28°C to 42°C (3-17°C above ambient) in 1°C increments (**Figure 1F**). After a rapid approach and equilibration at the designated set point, each well maintained its temperature with high accuracy (average error of 0.04°C from set point) and precision (average SD of 0.05°C within individual trace) over 15 hrs of operation (**Figure 1F**). Moreover, temperatures could be dynamically toggled between multiple setpoints, returning to the desired temperature with high fidelity (at steady state: average error of 0.07°C, average within-trace SD = 0.06°C) (**Figure 1G**).

### Characterizing speed and accuracy of thermoPlate heating

We next characterized key operational features of thermoPlate heating, including its range, accuracy, heating/cooling speed, and overshoot. We devised an experiment where the thermoPlate would sequentially heat each well from 3°C above ambient temperature to a prespecified higher temperature for 20 minutes, followed by cooling back to 3°C above ambient. Note that because the thermoPlate has no active cooling, in most experiments the baseline (lowest) experimental temperature should be set to higher than ambient to allow rapid cooling to—and maintenance of—this desired low temperature. **Figure 2A** demonstrates the above described experiment for heating through ΔT = 10°C. All wells experienced this heat pulse with high consistency between wells (**Figures 2A,B**). From this type of experiment, we could systematically quantify error (difference between the temperature and the set point after 20 minutes), rise/fall time (the time until temperature reached 99% of the target temperature), and overshoot/undershoot (difference between the maximum/minimum and setpoint).

**Figure 2.**
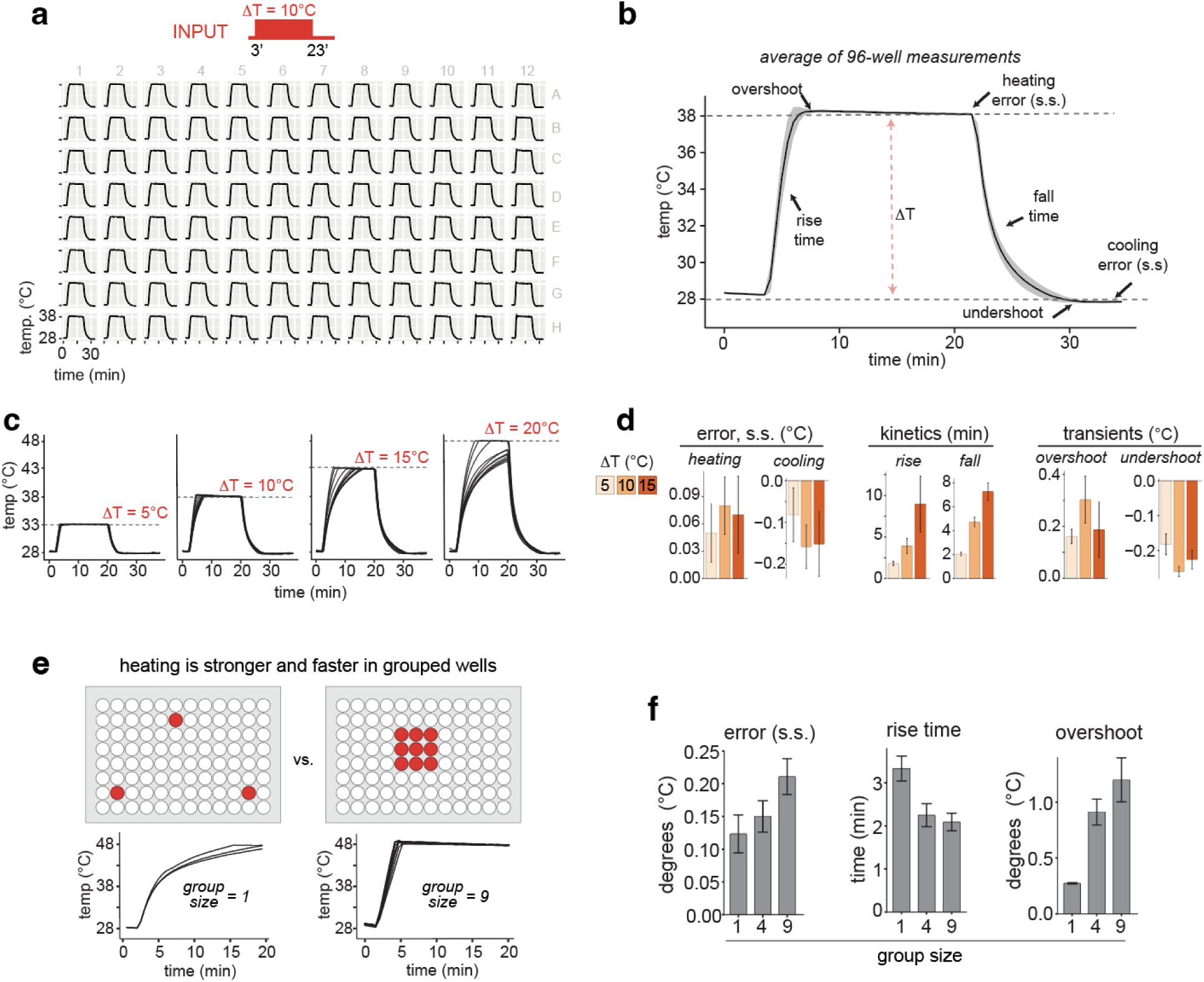
Characterization of thermoPlate heating and cooling. a) Temperature profile of each well heated with a 20 min pulse of 10°C. Wells were heated sequentially. b) The mean of all 96 traces from (a) labeled with key metrics of thermoPlate performance. Heating error = difference between the measured and set point temperature at 20 minutes. Rise time = duration at which temperature reached 0.5°C of the desired temperature. Overshoot = difference between maximum measured and set point temperature during heating. Cooling error, fall time, and undershoot represent the same parameters during the cooling phase. Error ribbons = 1 SD. s.s. = steady state. c) Heating each of 12 wells through 5, 10, 15, or 20°C for 20 minutes and then cooling to 28°C (ambient = 25°C). d) Mean of each performance metric from (c). Error bars represent = SD. e) Heating amplitude and speed can be increased by heating groups of wells. Wells were programmed to heat by ΔT = 20°C as either single wells or a group of 9 wells. f) Performance metrics of heating as a function of well grouping. Wells were heated by ΔT = 10°C. Data represents mean +/-SD from 3-9 wells.

To quantify performance as a function of heating magnitude, we chose 11 wells distributed across the plate and repeated the above experiment with heating of either 5, 10, 15, or 20°C above ambient, with each well heated sequentially (**Figure 2C**). While all wells reached their target temperature for up to ΔT = 15°C, only some wells could reach ΔT = 20°C within 20 minutes. The rate at which a well approached ΔT = 20°C was highly dependent on its position on the plate, with corner wells approaching 20°C more rapidly than center wells **(Figure S3).**

This variability in heating speed likely arises because of differences in heat diffusion in different parts of the plate, with higher numbers of neighbor wells acting as a larger heatsink that limits heating. Rise/fall times were 2 min for ΔT = 5°C and 4 min for ΔT = 10°C, with increasing variability between wells with increasing ΔT (**Figure 2D**). However, error (0.05-0.15°C) and overshoot/undershoot (0.1-0.3°C) were minimal and did not correlate with ΔT.

Although heat diffusion limited the heating range of individual wells, it could also be harnessed to boost heating within groups of wells. We tested this concept by heating either individual wells or groups of 4 or 9 wells to ΔT = 20°C. Although, as before, individual wells struggled to reach this level of heating, wells in a 3x3 matrix rapidly equilibrated to this high set point (**Figure 2E**). Grouped wells achieved a faster approach to high temperatures at the expense of small increases in error (0.15-0.2°C) and overshoot (∼1°C) (**Figure 2F**). In sum, the thermoPlate can reliably and rapidly achieve heating of up to ΔT = 15°C in individual wells and can also be programmed to achieve ΔT >20°C with appropriate arrangement of the heated wells.

### Modeling heat diffusion in the thermoPlate

Heat diffusion constrains the heating protocols that can be run simultaneously because it places limits on the temperature differential between two neighboring wells. For example, a well heated to 15°C above ambient will diffusively heat its neighboring wells by ∼10°C, placing a lower limit on the possible temperatures of those neighbors. As the number of wells in use increases, it becomes increasingly challenging to predict whether a desired protocol is achievable, particularly when dynamic heating protocols are desired.

We thus created a model of heat diffusion in 96-well plates to allow users to rapidly assess the feasibility of particular heating programs. We implemented an explicit finite difference approximation of a modified heat equation. This model simulates dynamic heat stimuli and calculates the rate of heat spread through each well of the plate over time. Unlike the classical heat equation, our model has 6 (vs 1) parameters that describe the rate of heat spread. These additional parameters account for observed differences in heat spread across different contexts, for example in edge wells vs center wells (**Figure 3A**). These parameters also accounted for small asymmetries around heated wells (left vs right, top vs bottom) likely due to asymmetric positioning of the electrical components.

**Figure 3.**
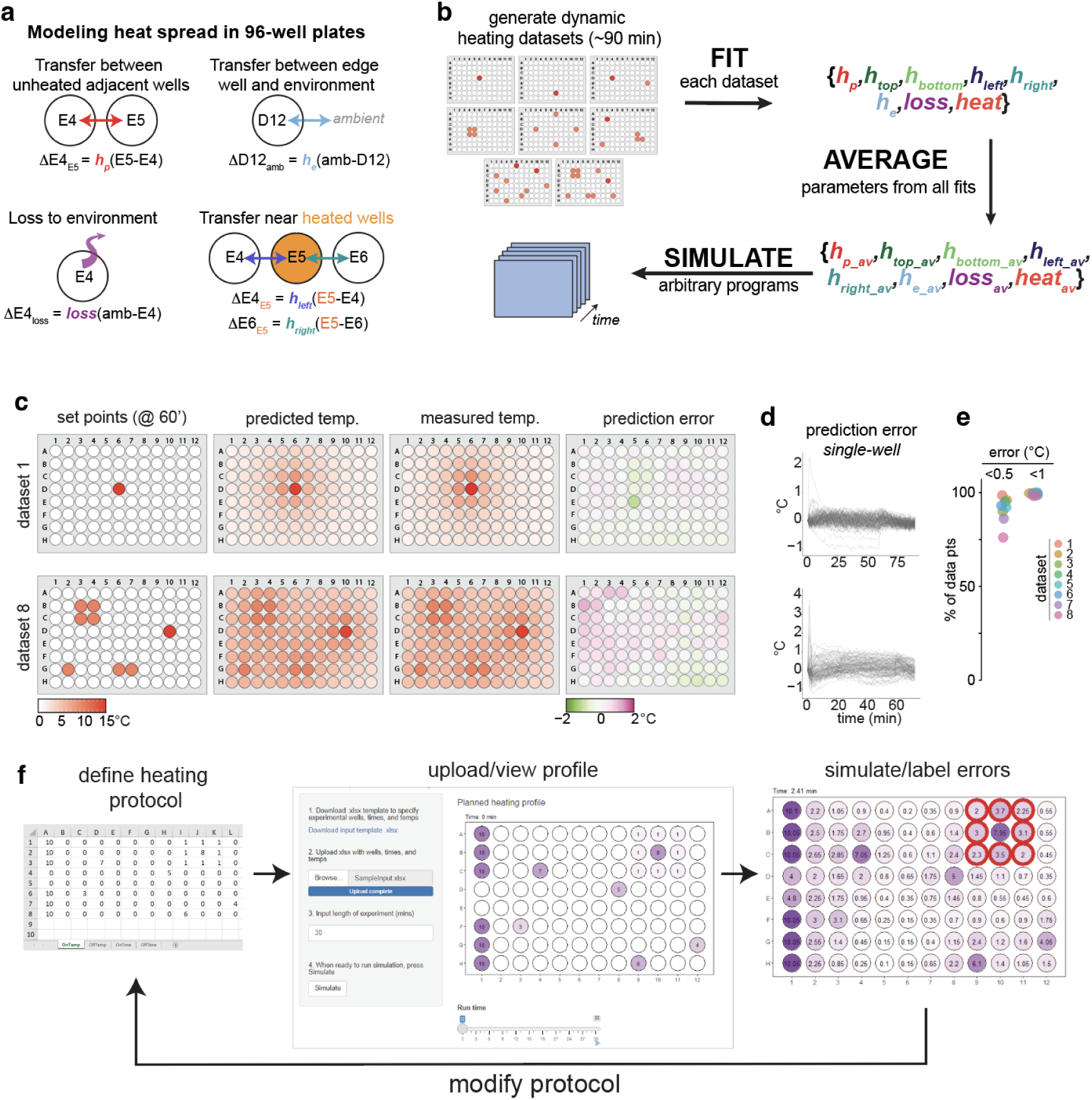
Mathematical modeling predicts the temperature of all wells in a sample plate. a) Diagram of parameters used to model heat diffusion in a 96-well plate that is heated by a thermoPlate. b) Strategy for parameter fitting. 8 parameter sets were fit to 8 separate experiments that spanned a range of wells, temperature, and heating dynamics. Parameters were then averaged across the parameter sets. c) Comparisons of set points, model predictions, actual temperatures, and error between predictions and model for the 60 min time points of two heating datasets. d) Traces of error (predicted-measured) within each well over the entire duration of two datasets. Each trace represents the error within a single well. e) Summaries of error for each well and time point of each data set. f) A Shiny App GUI allows users to upload heating profiles, review the profiles visually, and simulate heating and heat diffusion during a run. The simulation shows the predicted temperature of all wells and indicates whether any prespecified temperature profiles are unachievable due to thermal diffusion from neighboring wells (red circles in right panel). The user can then adjust and simulate the adjusted protocol until no errors are found.

To fit the model, we generated eight sets of training data where different sets of wells (1-12 wells) underwent dynamic heating/cooling programs, with temperature changes of 1-15°C over durations of up to 90 min (**Figure 3B**). These training experiments were designed with increasing levels of complexity, from simple heating of one well (Dataset 1) to more complex, dynamic heating of 12 wells arranged both individually and in groups (Dataset 8)(**Figure S4**).

The model was parameterized for each training set by minimizing the root mean squared error (RMSE). These 8 parameter sets were then averaged to create a final set of parameters that describe thermoPlate behavior. We validated the model using Leave-One-Out-Cross-Validation, wherein fitted parameters from 7 training sets were averaged and used to predict the 8th dataset, resulting in a mean average error of 0.23°C across all wells and times. Comparing the predictions of the full model vs the data across all data sets, the average error over an experimental run was 0.22°C (**Figure 3C-E**), with error falling within 1°C for > 99% of timepoints and within 0.5°C for > 75% of timepoints (**Figure 3E**), showing close agreement between model and data. Transient spikes in error (1-2°C) could on occasion be observed near wells that experienced a sudden large temperature change (**Figure 3D**).

The predictive power of this model allows users to rapidly test whether particular heating patterns and spatial arrangements are permissible. For this purpose, we created an online interface where users can upload the desired heating protocol and easily simulate and visualize heat generation and spread (**Figure 3F**). The visualization also highlights impermissible temperature set points, allowing the user to modify their experimental design accordingly.

### Controlling and characterizing thermogenetic phase separation

We applied the thermoPlate to probe and observe thermally-controlled biochemistry in mammalian cells. We first confirmed that the thermoPlate was fully compatible with mammalian cell culture, with no observed impact on proliferation or survival of HEK 293T cells after even 48 hr of direct contact with the thermoPlate (**Figure S5**). Measurement of evaporation revealed that, while significant evaporation could be observed in edge wells under prolonged heating (24 hr at >10°C), evaporation could be prevented by adding a thin layer of paraffin oil over each well (**Figure S6**).

We first leveraged rapid and dynamic thermal control to characterize phase separation of an elastin-like polypeptide (ELP_53_), which forms protein condensates in mammalian cells above a critical temperature (**Figure 4A,B**)^11^. ELP condensates form rapidly (∼minutes) and over a small temperature range (∼1-2°C)^12,13^. We used the thermoPlate coupled with live-cell confocal microscopy to observe ELP_53_ condensation in response to increased temperature between 28-39°C, at 1°C resolution (**Figure 4C**). ELPs condensed rapidly and as a function of temperature, with measurable condensation above 31°C and a ∼linear increase in magnitude between 33-38°C (**Figure 4C,D**). We next quantified the speed of ELP formation kinetics, finding that ELP condensation increased within 1 min of heating and reached steady-state within 10 min (**Figure 4E**). Notably, the thermoPlate achieved its target temperatures in <2 minutes, sufficiently fast to observe the slightly slower ELP condensation. The fast and dynamic responses of both the thermoPlate and ELPs allowed us to reversibly toggle ELP phase separation over 45 cycles of heating and cooling (4 min 35°C, 4 min 29°C). ELP condensation and dissolution remained consistent over all cycles, with no decrease in dynamic range, suggesting that such stimulation could be maintained indefinitely (**Figure 4F, Movie S1**).

**Figure 4.**
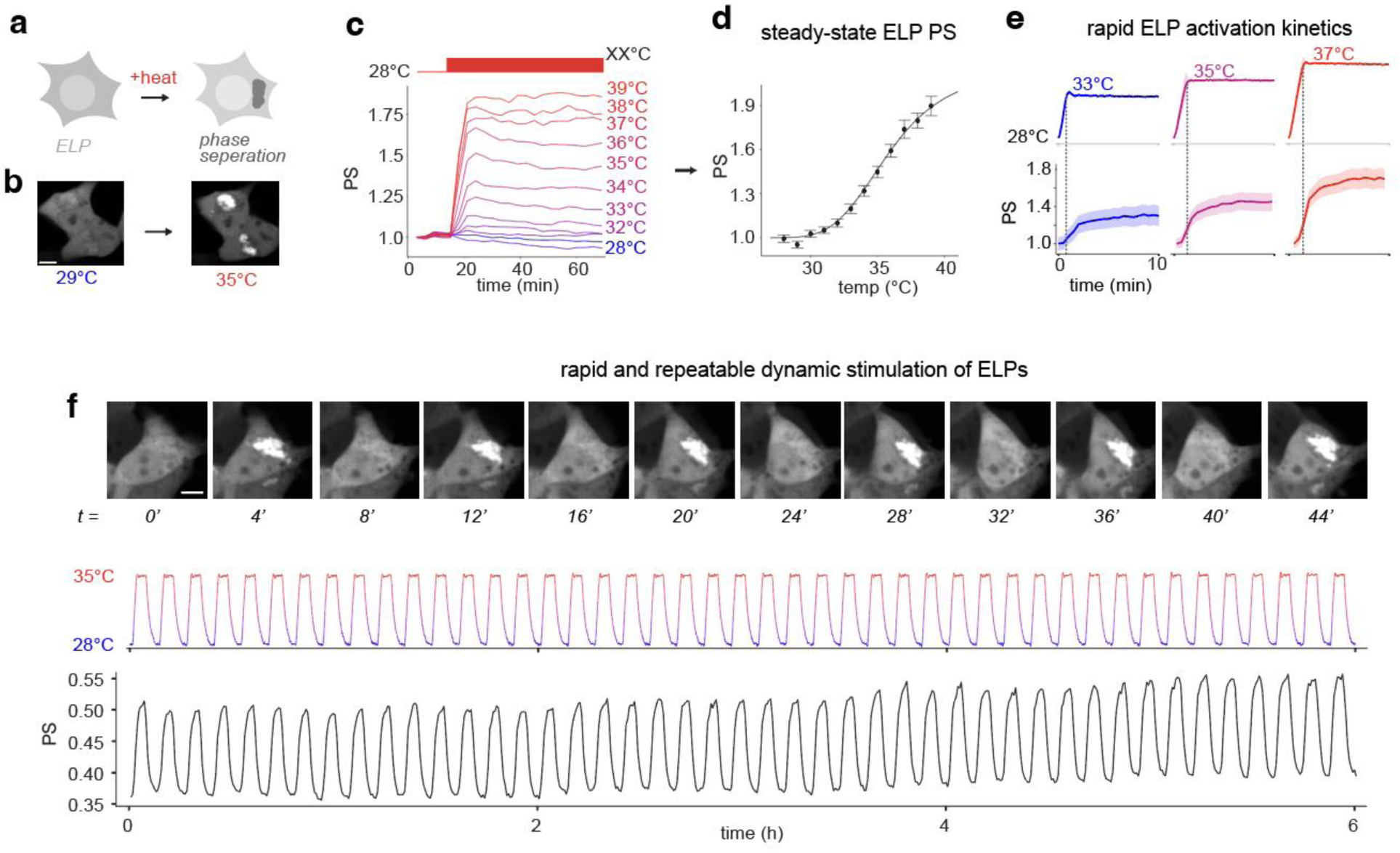
The thermoPlate allows rapid characterization of the phase separation of an elastin-like polypeptide (ELP_53_). a) Diagram of ELP phase separation (PS) upon elevated temperature. b) Representative image of ELP phase separation when transition from 29°C to 35°C in HEK 293T cell lines. c) Cells expressing ELP-GFP were exposed to various temperatures for 1 hour while undergoing simultaneous confocal microscopy. Each trace represents the mean of ∼500 cells. PS values are normalized to t = 0. See methods for PS quantification. d) The equilibrium PS at each temperature from (c) plotted vs temperature. Each point represents the mean of ∼500 cells +/-1 SEM. PS is normalized to values at 28°C. e) Quantification of the kinetics of ELP PS at 3 temperatures. Traces represent the mean of ∼500 cells +/-1 SEM. PS values are normalized to t = 0. f) Rapid and dynamic control of ELP PS can be maintained indefinitely. Data show 45 cycles of 4 minutes of heating to 35°C followed by 4 minutes at 28°C. The temperature trace represents the mean of 3 wells. PS traces represent the mean of ∼1000 cells.

### Characterizing dynamics and memory of heat stress in mammalian cells

We next applied the thermoPlate to examine native thermal responses of mammalian cells. We focused on the dynamics of stress granule formation in response to heat shock. Stress granules (SGs) are condensates of protein and RNA that form in the cytosol in response to stressors including heat, hypoxia, and oxidation^14^. While the composition and function of SGs in response to heat stress have been studied extensively^15–20^, their dynamics are relatively less explored. To observe stress granules, we stably expressed a fluorescent fusion of the SG scaffold G3BP1-mCherry in HeLa cells, and we imaged G3BP1 condensation as a function of time and temperature.

We first asked at what temperature SGs could be observed by exposing cells to constant heat between 41 to 47°C (**Figure 5A**). We found a highly non-linear relationship between temperature and the magnitude of stress-granule formation. Although no SGs were observed at 41°C, SG rapidly formed at 42°C and increased in magnitude at 43°C and 44°C. Surprisingly, maximal SG intensity decreased with progressively further increases in temperature (**Figure 5B-D**). Although cells remained morphologically unperturbed with SGs at 42°C and 43°C over at least 8 hrs, cells heated with 44°C lost cytoplasmic volume and became immobile, with higher temperatures leading to faster onset of this phenotype (**Figure 5C, Movie S2).** In all heating conditions, SGs formed rapidly within the first 5 min.

**Figure 5.**
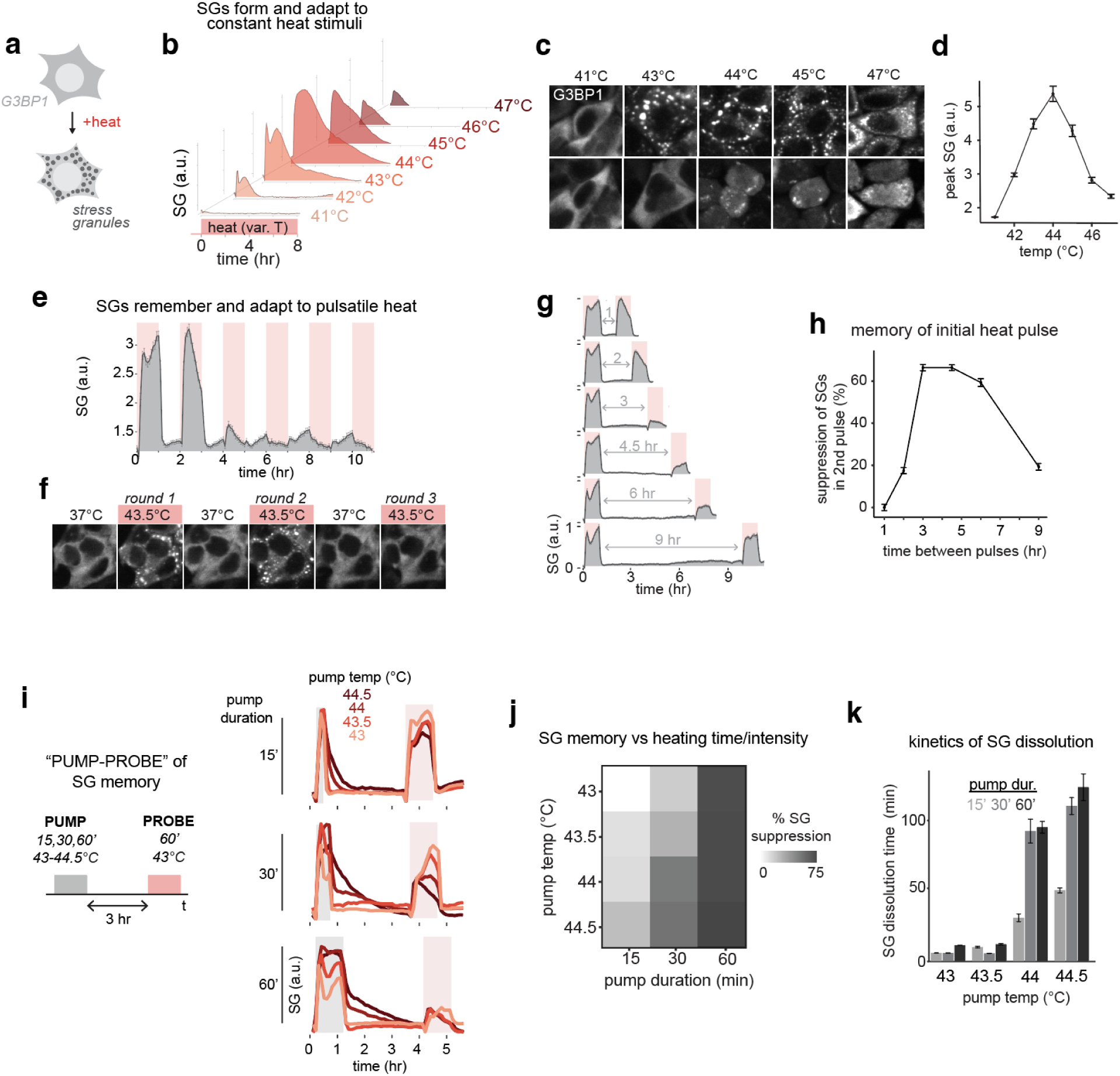
The thermoPlate allows visualization of mammalian heat shock response with high resolution in temperature and time. a) Heat induced stress granule (SG) formation was observed by measuring condensation of G3BP1-mCh in response to time varying temperature changes in HeLa cells. b) SG formation in response to constant heating. Each trace represents mean SG formation from two wells. c) Representative images of experiment in (b). d) Maxima of SG formation across all temperatures tested. e) SG response dynamics in response to pulsatile stimuli. Red bars indicate heating intervals (1 hr at 43.5°C), which were separated by 1 hr intervals at 37°C. f) Interrogating temporal properties of heat shock memories that suppress stress granule formation. Cells were heated for 1 hr intervals (43.5°C) separated by variable intervals at 37°C, and SG formation on the second pulse was measured. h) The degree of suppression of SGs on the second pulse from (g) is quantified as a function of the interval between heat pulses. i) A “pump-probe” experiment to interrogate the effects of temperature and duration of heat shock on SG memory. The temperature and duration of the “pump” heat pulse were varied, and the magnitude of SG formation on a second heat pulse (1 hr, 43.5°C) was measured. j) Quantification of SG memory (suppression during the probe pulse) as a function of pump temperature and duration from experiment in (i). k) Quantification of kinetics of SG dissolution after the pump pulse. Dissociation time was calculated as the time at which SG magnitude dropped 90% from its peak value before adaptation. Data represents mean +/-SD of four replicates.

Notably, SG formation was transient across all conditions despite constant heating, and in most cells G3BP1 returned to a diffuse state within several hours of the onset of heat shock (**Figure 5B**). We asked if SG adaptation required constant heat stress or alternatively could form in response to short heat pulses. We examined SG formation in response to periodic 1-hr pulses of heat stress (43.5°C), a pulse duration shorter than the time scale of SG adaptation (**Figure 5B**). Although the SGs formed and dissolved in sync with the first two heating and cooling pulses, SGs largely did not form during the 3^rd^ and subsequent pulses, suggesting that cells continued adapting to heat stress despite heat not being applied for the entire duration of adaptation (**Figure 5E,F, Movie S3**). Indeed, even though the second heat pulse yielded strong SG formation, SG magnitude began to decrease during this second window, indicating that the first heat pulse influenced SG behavior during the second heat pulse (**Figure 5E**). Collectively, these results show that cells store memories of prior heat exposure that shape future responses to repeated stress.

We next characterized the timescales of this memory formation. We programmed the thermoPlate to apply two pulses of heat stress (43.5°C) separated by varying periods of relaxation at 37°C (**Figure 5G**), and we quantified SG formation during the second pulse. SG formation depended on the interval between pulses in a biphasic manner. Although SGs were prominent during the second pulse after a 1 or 2 hr interval, they were not observed with intervals of 3 and 4.5 hrs. At still longer intervals, SG formation began to reappear on the second pulse. With a 9 hr interval, SG formation during the 2nd pulse resembled that of the first pulse (**Figure 5G,H**). Thus, cellular memory of prior stress was maximal for 3-6 hrs after which the memory began to fade and cells again responded to heat shock as on the first exposure.

Finally, we wondered whether memories of prior heat shock depended on the duration or the temperature of heat shock, or both. We designed a “pump-probe experiment” where we delivered a “pump” of heat stress with variable duration (15, 30, 60 min) and temperature (43-44.5°C), returned to 37°C for 3 hr, and then “probed” SG formation with a subsequent 1 hr of heat shock (43.5°C) (**Figure 5I**). Indeed, memory (suppression of SG formation on the second pulse) depended on both time and temperature of the initial heat shock pump. Suppression of SGs was stronger with increasing temperatures, and the magnitude of suppression increased with increasing pump duration. Notably, even a short 15 min pump was sufficient for measurable SG suppression at higher temperatures (**Figure 5I,J**). Additionally, the rate of dissolution of SGs after removal of the pump pulse depended on the intensity and duration of heat stress, with slower dissociation in response to longer and more intense heating (**Figure 5I**), indicating a second form of stress memory. This effect could only be observed at or above 44°C heating and could be observed even after a brief 15 min pump duration (**Figure 5K**). In sum, our results indicate multiple mechanisms by which cells can store memories of prior heat stress and demonstrate the utility of the thermoPlate to explore these stress responses efficiently with high precision, throughput, and temporal resolution.

## Discussion

The thermoPlate allows modulation of the temperature of biological samples in a multiplexed and small format that is compatible with both long term cell culture and live-cell microscopy. After calibration, the thermoPlate achieved its desired set point temperatures rapidly and accurately, with minimal variation between wells, allowing profiling of the response to thermal dynamics with high resolution in temperature and in time. The device is open source and can be assembled by non-specialists in several hours by following the assembly protocols associated with this manuscript, for a total cost of ∼$300 in parts. The design files can also be modified to adapt the device for user-specific applications as needed, for example for use in larger well-plate formats.

Heat diffusion constrains the types of heating protocols that can be run simultaneously. Nevertheless, these constraints need not limit throughput. For example, if a sustained temperature sweep is desired, a temperature gradient can be specified across the plate, and all 96-wells can be used simultaneously. For more complex and dynamic patterns, samples must be placed more strategically to allow the samples to heat and cool appropriately. While this is straightforward for a few samples, it becomes more challenging with each additional heated well. For this reason, we developed a model of thermal diffusion that accurately predicts heating dynamics of each well for arbitrary protocols. The user can rapidly simulate a desired protocol *in silico* and observe whether the intended thermal patterns are permitted within the constraints of thermal diffusion. If they are not, the user adjusts the inputs, simulates, and iterates until a suitable arrangement is reached. We also introduce a web-based GUI for easy access to this model.

The thermoPlate also harnesses thermal diffusion to passively cool the samples. The simplicity of passive cooling is balanced by two additional constraints: 1) potentially slow cooling kinetics and 2) inability to maintain certain wells at ambient temperatures while heating other wells, since over time all wells will increase temperature by a few degrees as heat spreads throughout the plate faster than it diffuses out to the environment. To address both concerns, we maintained the environment at least ∼3°C cooler than the lowest experimental temperature. This ensures that low experimental temperatures can be maintained and also results in faster cooling to this low level. With this approach, heating and cooling occur on comparable ∼minutes timescales (**Figure 2D)**.

The thermoPlate enables rapid characterization of new thermogenetic tools, where temperature acts as an inducer for genetic or protein activity^6–8^. We demonstrated this ability by systematically measuring the steady-state, kinetics, and dynamics of ELP_53_, an elastin-like peptide that forms condensates above a critical temperature. Although ELP phase separation is rapid, the fast heating of the thermoPlate was able to capture rapid ELP condensation and its slower approach to equilibrium, showing kinetics that are consistent with prior reports of ELP condensation in cells^11,12^. ELP condensation could also be toggled indefinitely with fast minutes-scale ON- and OFF-kinetics. Such rapid clustering matches the timescales of even the fastest optogenetic clustering approaches^21^. In a separate report, the thermoPlate was used to systematically characterize the thermogenetic Melt protein, further demonstrating the potential of the thermoPlate for thorough characterization of temperature-sensitive proteins^22^.

The thermoPlate also empowers studies of native cellular responses to temperature, which we demonstrated by exploring the dynamics of the stress granule (SG) formation in response to time-varying heat-shock temperatures. We found a rich set of dynamic features of SG formation that respond to both the time and temperature of heat stress in a non-linear manner. Rather than a binary response, SG formation differed with each additional degree of heating above 41°C, with a maximal peak amplitude at 44°C. Notably, SGs showed adaptation to heat stress, such that G3BP1 puncta spontaneously dissolved despite persistent heating. Adaptation of SGs has been previously reported in the context of oxidative stress^23^ and viral infection^24^. Such adaptation is consistent with negative feedback, including through transcription of heat shock proteins (HSPs)^25–29^ and of suppressors of the integrated stress response including GADD34^24,30,31^. Nevertheless, to our knowledge the adaptation of SGs to chronic heat stress has not been previously reported, demonstrating the utility of simultaneous thermal control and live-cell imaging.

Heat stress resulted in the formation of at least two types of biochemical memories. A 1 hr pulse of stress was sufficient to suppress SG formation upon subsequent stress, with maximal suppression observed 3-6 hrs after the initial stress pulse. These kinetics are in line with a recent finding that 1 hr of heat stress induced suppressed activation of the integrated stress response after a second heat pulse^24^. However, this prior work did not observe the status of SG formation during this second pulse. Notably, our measurements also revealed SG suppression after only 15 and 30 min of initial heat shock. With these shorter durations, the strength of SG suppression increased with increasing temperature of prior heat shock. A second form of biochemical memory appeared in the kinetics of SG dissolution after stress removal. This memory was observed immediately after removal of heat stress but only above 44°C, consistent with fast post-translational mechanisms like protein clustering distinct from SG formation^32^. While further work will be required to identify the specific mechanisms that underlie our observations, our results highlight the discovery potential of instrumentation for easy, precise, and dynamic control of temperature under a microscope.

## Supporting information

Repository including usage manuals

Files for running the thermoPlate GUI

## Acknowledgements

The authors thank Dr. James Shorter (Penn) and Dr. Steven Boeynaems (Baylor) for helpful discussions. This work was supported by the National Science Foundation (CAREER CBET 2145699 to L.J.B., GRFP to W.B.), the National Institutes of Health (R35GM138211 to L.J.B), and the Penn Center for Precision Engineering for Health (CPE4H). Cell sorting was performed on a BD FACSAria Fusion that was obtained through an NIH S10 grant (S10OD026986).

## Author Contributions

W.B. and L.J.B conceived study. W.B. P.I., L.J.B. designed hardware structure and component layout. W.B. and P.I. constructed hardware, performed experiments, and analyzed data. T.M. and P.I. developed the model of thermal diffusion. Z.H. generated the ELP_53_ construct. L.J.B. supervised the work. W.B., T.M., and L.J.B. drafted manuscript and made figures. All authors contributed to editing.

## METHODS

### PID Implementation

PID (Proportional, Integral, Derivative) control was implemented using a discrete form of the standard PID algorithm. PID constants were generated empirically by testing several combinations. In brief, the Arduino collects the temperature of every well every ∼2 seconds. These values are used to calculate the current error (proportional), change in error from the previous reading per unit time (derivative), and integral (time weighted sum of all past error). The algorithm applies the PID constants to each error value and calculates a new duty ratio to apply to the corresponding heater. For more detailed information on programming architecture, see the **Code** section of the **Repository**.

### thermoPlate Calibration

Calibration was performed by placing a thermoPlate on a 96 well plate containing PBS. This assembly was equilibrated in a standard mammalian cell culture incubator at ∼25°C. Two digital thermometers (RC-4 Elitech Digital Temperature Data Logger) were placed inside two 15mL conical tubes filled with water and sealed with parafilm (to prevent evaporation). Both conical tube/thermometer were placed next to the thermoPlate, and a plastic box was placed over the thermoPlate and thermometers to avoid variation due to airflow. The average of the two digital thermometers was recorded as the true temperature. 5 temperature readings were recorded from each well. The average of those 5 readings was assigned as the final temperature reading of that well. The incubator temperature was then increased to 30°C and allowed to equilibrate for 2 hours. Readings from the thermoPlate and the digital thermometers were recorded, and this procedure was repeated in 5°C steps until 45°C. For each well, the temperature readings of the thermoPlate vs digital thermometers was then plotted, and the slope and y-intercept of the linear fit were extracted. These values were used to calibrate each raw temperature reading during thermoPlate operation. See **Manual** for more details on thermoPlate operation.

### thermoPlate Modeling

A model of heat transfer within a thermoPlate-heated 96-well plate was made in MATLAB to simulate the temperature of each well given a desired heating input. Well temperatures were simulated through time. At each timestep, the instantaneous heat derivative of each well was calculated and multiplied by the simulation timestep to obtain the well’s temperature at the next timestep. The model uses 8 parameters. **H_p_** describes heat transfer between neighboring unheated wells. Around heated wells, **H_left_**, **H_right_**, **H_top_**, and **H_bottom_** describe heat transfer between the heated well and its neighbors to the left, right, top, and bottom, respectively. This was done to capture the consistent asymmetries seen in temperature around wells that are actively heated. **H_e_** describes heat transfer from an edge well to the environment, ***loss*** describes a consistent loss of heat from each well to the environment. Finally, ***heat*** is a constant temperature increment that is added to each heated well during a simulation step where the well is below its set temperature. When a heated well is at or above its setpoint, this ***heat*** is not added for that step. Example calculations of the change in temperature of wells over a single time step are shown in **Figure S7**. Parameters were obtained by fitting the model to eight real datasets and averaging resultant 8 parameter sets to obtain generalizable parameters. Fitting was done by minimizing an error function, defined as the difference between the simulation and the real data summed across every well and timepoint. Minimization was done using the MATLAB function fminsearch.

### Cell Culture

HeLa cells were maintained in 10% fetal bovine serum (FBS) and 1% penicillin/streptomycin (P/S) in DMEM. Cell lines were not verified after purchase. Cells were not cultured in proximity to commonly misidentified cell lines.

### Plasmid design and assembly

Constructs for stable transduction and transient transfection were cloned into the pHR lentiviral backbone with an SFFV promoter driving the gene of interest. The pHR backbone was linearized using MluI and NotI restriction sites. G3BP1 (Ophir Shalem Lab) and mCherry inserts were generated via PCR and cloned into the linearized viral backbone as a single fusion protein using HiFi cloning mix (NEB). H2B-iRFP nuclear marker was (Addgene Plasmid #90237). ELP_48_ was obtained from Addgene (Addgene Plasmid #68395). Sequencing of ELP_48_ showed it contained 44 ELP repeats. ELP_48_ was digested using NdeI and BamHI and ligated into a small backbone pTA for the ease of cloning. ELP_9_ was ordered as primers and inserted via HiFi into the pTA-ELP_48_ linearized with AgeI and BsmBI, generating pTA-ELP_53_. EGFP was then amplified and inserted via HiFi to pTA-ELP_53_, which was linearized using MluI and XcmI. ELP_53_-EGFP was then subcloned via HiFi to the viral backbone pHR digested with MluI and NotI.

### Plasmid transfection

HEK 293T cells were transfected using the calcium phosphate method, as follows: Per 1 mL of media of the cell culture to be transfected, 50 µL of 2x HeBS buffer, 1 µg of each DNA construct, and H_2_O up to 94 µL was mixed. 6 µL of 2.5mM CaCl_2_ was added after mixing of initial components, incubated for 1:45 minutes at room temperature, and added directly to cell culture.

### Lentiviral packaging and cell line generation

Lentivirus was packaged by cotransfecting the pHR transfer vector, pCMV-dR8.91 (Addgene, catalog number 12263), and pMD2.G (Addgene, catalog number 12259) into Lenti-X HEK293T. Briefly, cells were seeded one day prior to transfection at a concentration of 350,000 cells/mL in a 6-well plate. Plasmids were transfected using the calcium phosphate method. Media was removed one day post-transfection and replaced with fresh media. Two days post-transfection, media containing virus was collected and centrifuged at 800 x g for 3 minutes. The supernatant was passed through a 0.45 µm filter. 500 µL of filtered virus solution was added to 700,000 HeLa cells seeded in a 6-well plate. Cells were expanded over multiple passages, and successfully transduced cells were enriched through fluorescence activated cell sorting (BD FACS Aria Fusion).

### Preparation of cells for plate-based experiments

All experiments were carried out in Falcon™ 353219 Black 96 well plates. Briefly, wells were coated with 50 uL of MilliporeSigma™ Chemicon™ Human Plasma Fibronectin Purified Protein fibronectin solution diluted 100x in PBS and were incubated at 37 °C for 30 min. HeLa or HEK 293T cells were seeded in wells at a density of 60,000 cells/well in 100 µL and were spun down at 20 x g for 1 minute to ensure an even distribution of cells across the wells. For imaging ELPs, HEK 293T cells were transfected with GFP-ELP_53_ and H2B-iRFP 12-24 hours post-seeding in 96 well plates using the calcium phosphate method. Transfection mix/media was removed 24 hours post transfection and replaced with fresh media. Cells were imaged within 4-8 hours of this media change.

### Imaging

#### Live-cell imaging

Live-cell imaging was performed using a Nikon Ti2-E microscope equipped with a Yokagawa CSU-W1 spinning disk, 405/488/561/640 nm laser lines, an sCMOS camera (Photometrics), a motorized stage, and an environmental chamber (Okolabs). Cells expressing the construct of interest were imaged with a 20X air objective at variable temperatures and 5% CO_2_.

#### Imaging quantification

Images were processed using Cell Profiler ^33^. Cells were segmented using the H2B-iRFP imaging channel, and cytoplasm was identified using a 20 pixel ring around the nucleus. Nuclear and cytoplasmic fluorescence values were then exported and analyzed using R (https://cran.r-project.org/) and R-Studio (https://rstudio.com/). Data was processed and visualized using the tidyR ^46^ and ggplot2 ^47^ packages. Stress granules and ELP condensates were quantified by averaging the maximum pixel intensity of each cell within a frame and then averaging those values across multiple replicates.

## Supplementary Figures

**Figure S1.**
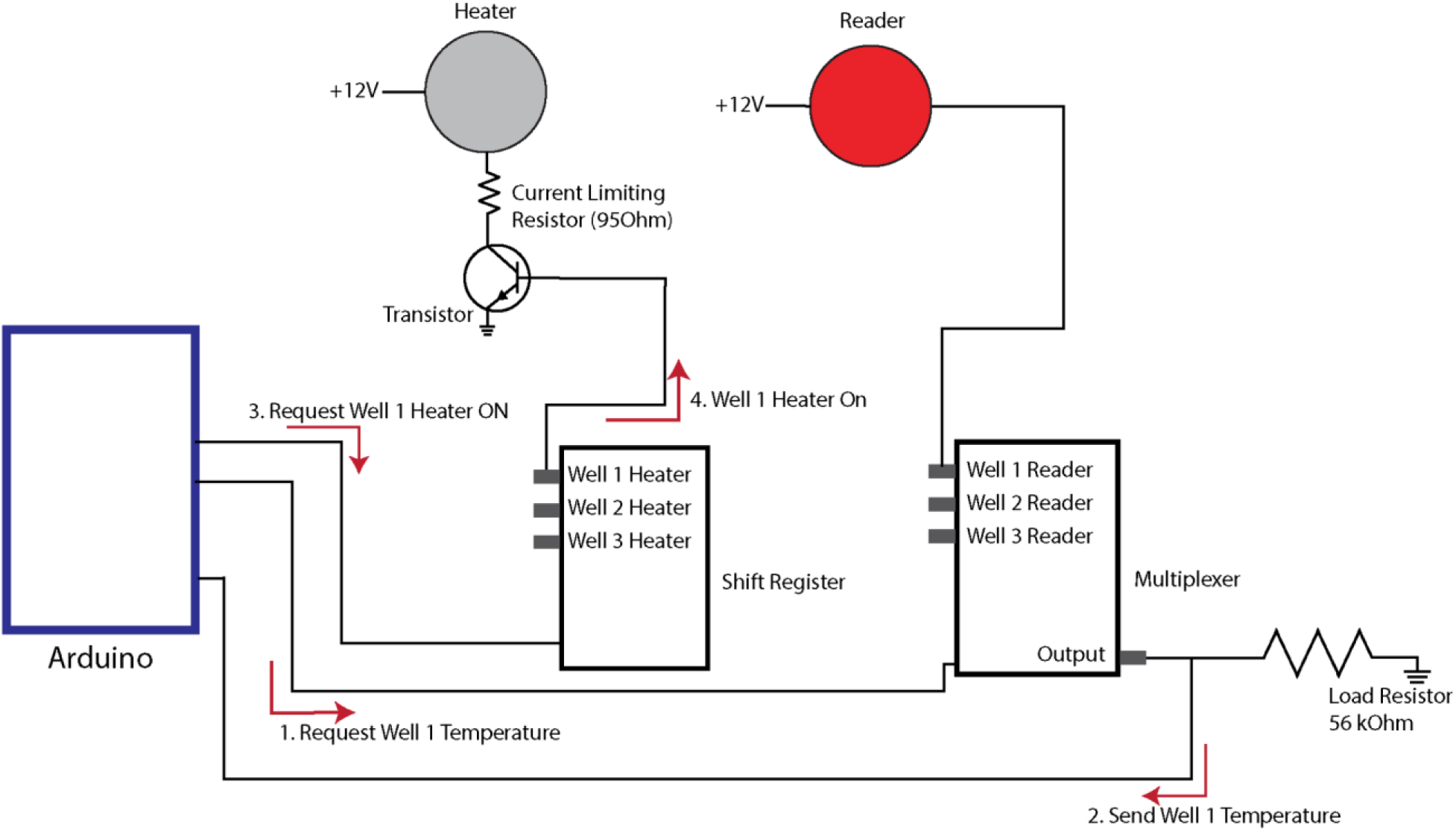
Wiring diagram of a single Heater/Reader thermistor pair. Each Heater/Reader pair is controlled via an Arduino microcontroller which coordinates the following actions: 1) The temperature of a well is requested by the Arduino from the multiplexer. 2) The multiplexer retrieves and returns the temperature of the well via the Reader. The Arduino calculates the necessary heating intensity (in the form of a duty cycle) via a PID algorithm. 3) The Arduino requests that the shift register accordingly adjusts the on/off state of the Heater. 4) The shift register will adjust the Heater accordingly. This process is repeated 96 times for each well of a 96-well plate.

**Figure S2.**
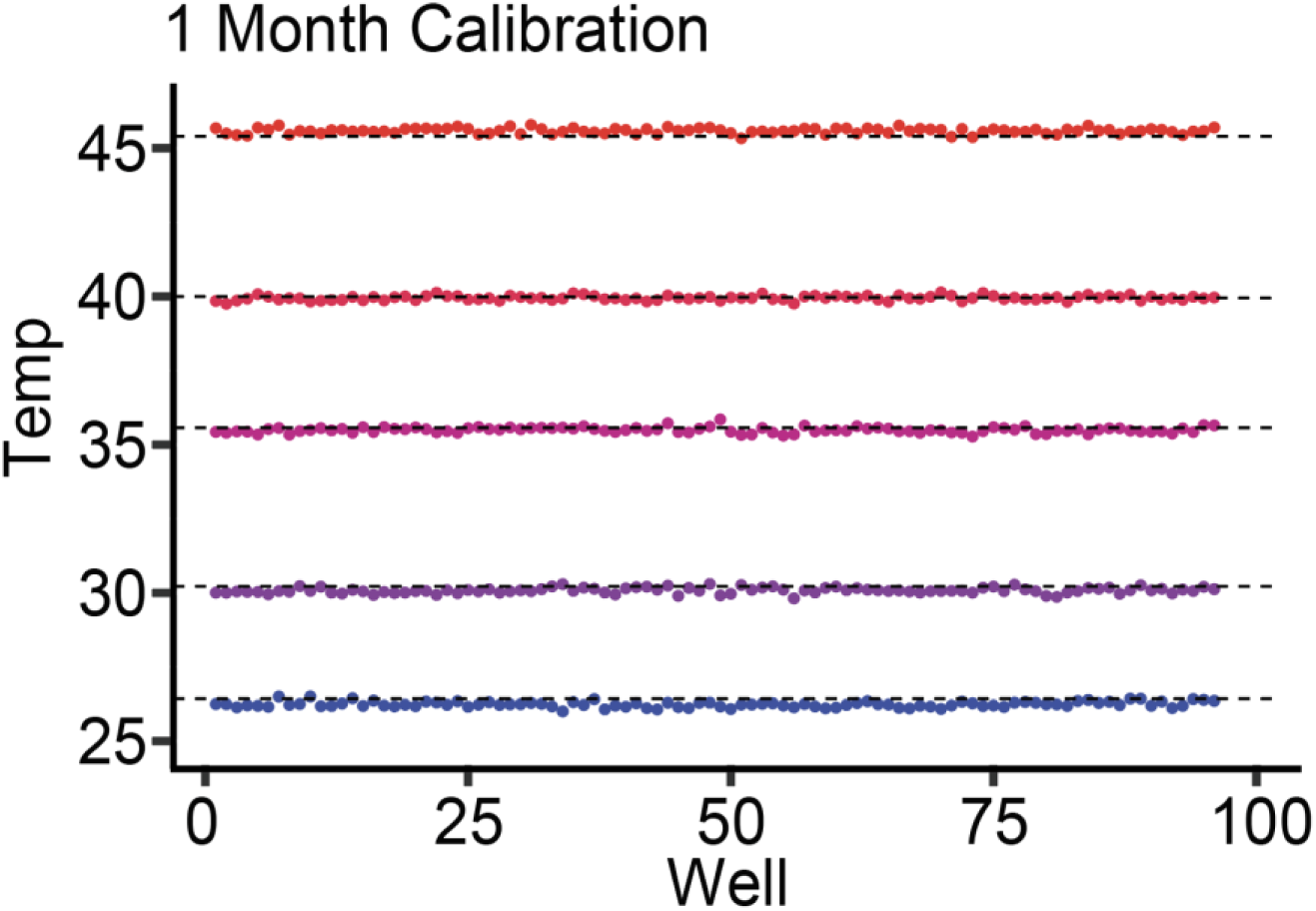
Calibration stability after 1 month of use. a) The same procedure performed in Figure 1e (raising the ambient temperature of an incubator in 5°C increments and recording thermoPlate readings at those temperatures) was repeated on the same thermo Plate 1 month after initial calibration, with temperature readings adjusted using the same calibration values produced in Figure 1e. Average reading error remained unchanged compared to prior calibration (<0.1°C) while average SD between wells increased marginally to 0.086°C, indicating calibration values are stable over time. Color and dashed lines represent the incubator temperature. See Methods for details on calibration.

**Figure S3.**
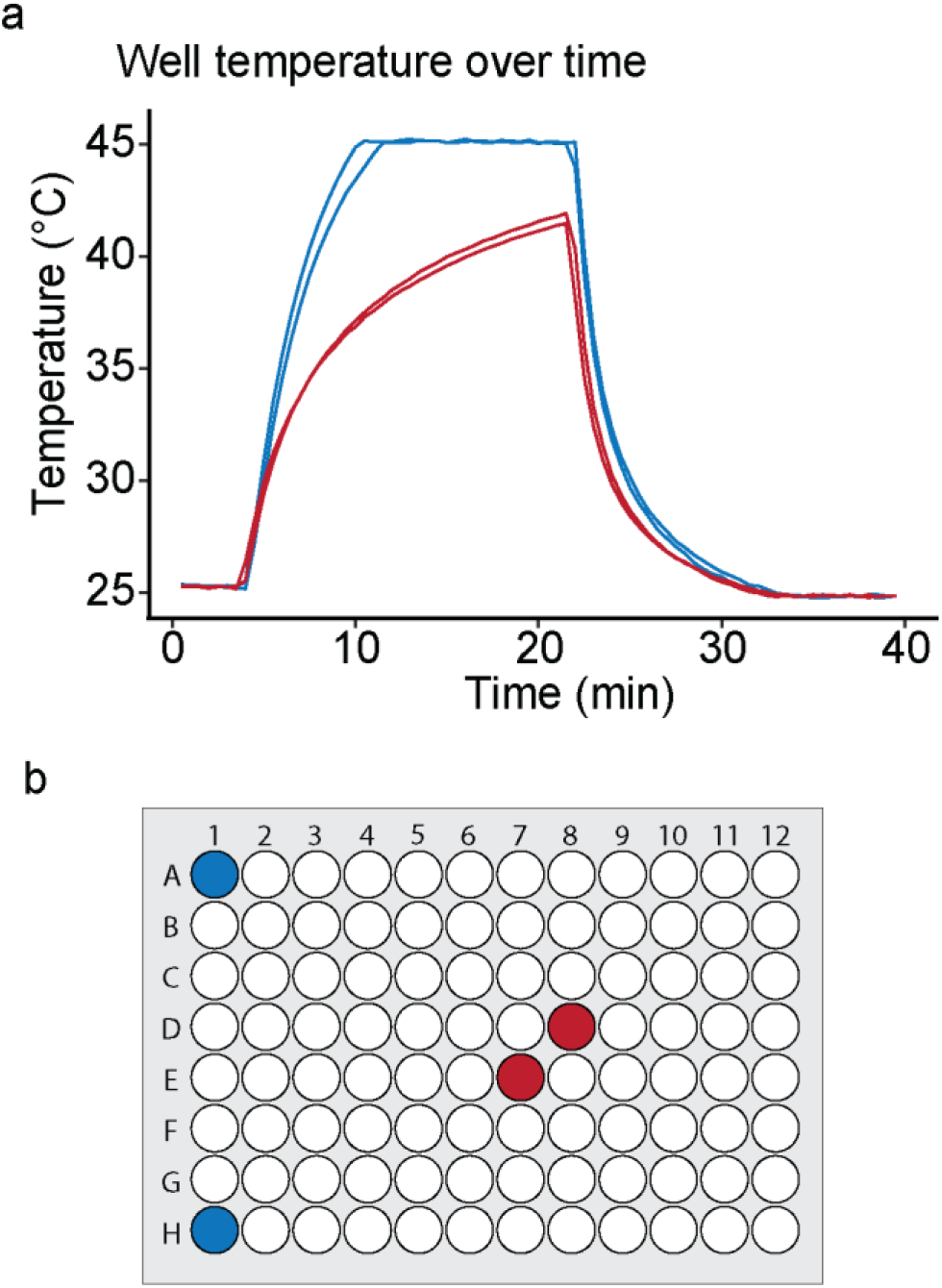
Edge wells heat faster and reach higher steady states. a) Traces are taken from Figure 2c, showing kinetics and steady state temperature from wells undergoing constant heating. Color represents the position on the plate (center wells labeled in red and edge wells labeled in blue) showing that edge wells are able to achieve faster and higher steady state heating than center wells. b) Diagram of the position on the plate of each trace found in (a).

**Figure S4.**
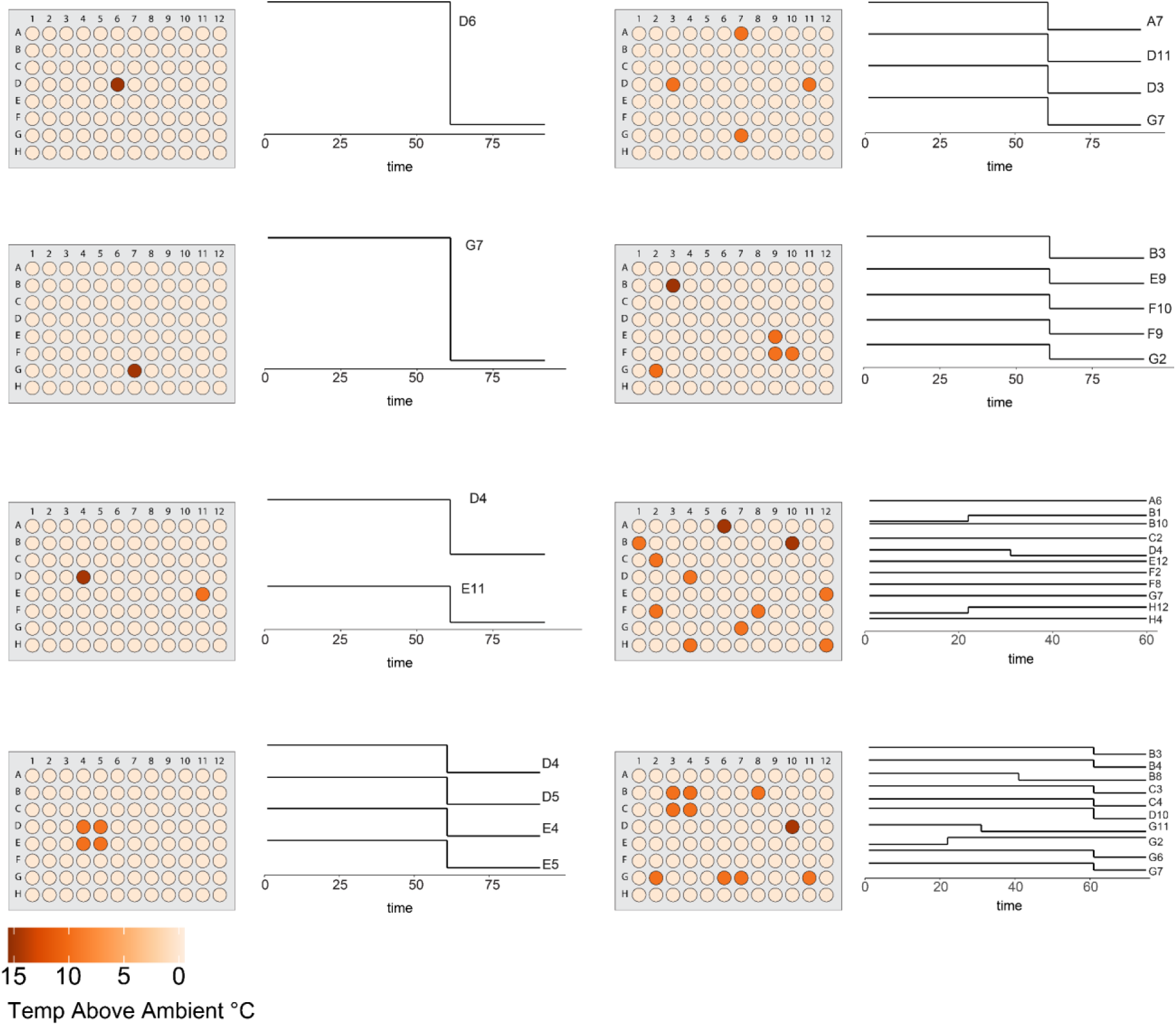
Visualization of heating profiles used to fit the thermoPlate model. a) Each diagram corresponds to one dataset used for model fitting with color representing the maximum set temperature of each well throughout the experiment. Traces represent the times at which specific wells either raise or lower their setpoint, normalized between max and min for that well.

**Figure S5.**
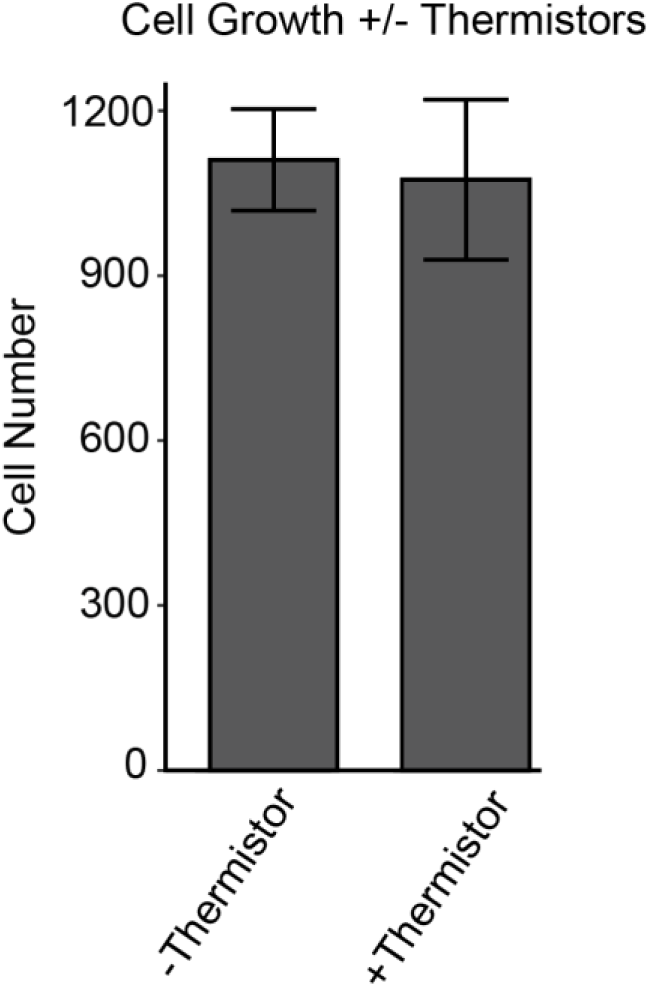
Cell growth quantification with and without exposure to the thermoPlate. a) Every well of a 96 well plate was seeded with HeLa cells and kept for two days in a standard cell culture incubator, with half of the wells exposed to thermoPlate heater/readers. Cells were then fixed, stained with DAPI, imaged, and nuclei were counted to quantify population size in each well. No significant difference in cell numbers was observed between the two conditions, indicating that cell growth and survival are not significantly altered by thermoPlate exposure. Each bar represents the mean +/-SD of N = 48 wells.

**Figure S6.**
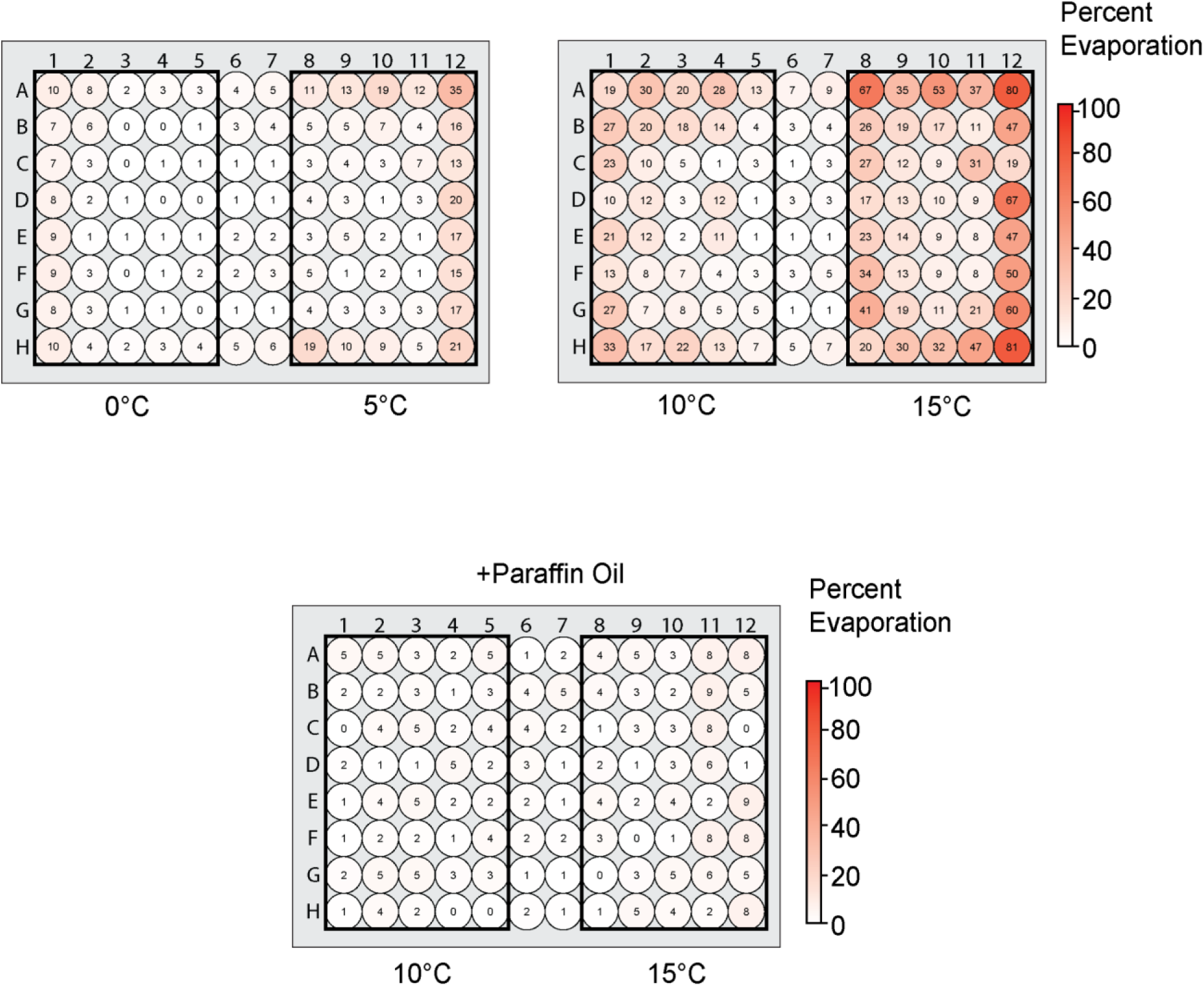
Evaporation during thermoPlate usage. a) A 96 well plate filled with 150uL of PBS was placed in a cell culture incubator and wells were set to heat by either 5 or 10°C for 24 hours. The volume of PBS remaining in the well was measured via a micropipette. Color represents the percent evaporation in each well. b) The same procedure as (a) was repeated heating wells by either 15 or 20°C. Major evaporation (>50%) was limited to wells set to 20°C above ambient. c) The same procedure as (b) was performed with a thin layer of 50uL of paraffin oil coating each well to prevent evaporation. Evaporation in all wells was <10% regardless of heating demonstrating that paraffin oil can be used to extend the duration of long-term heating.

**Figure S7.**
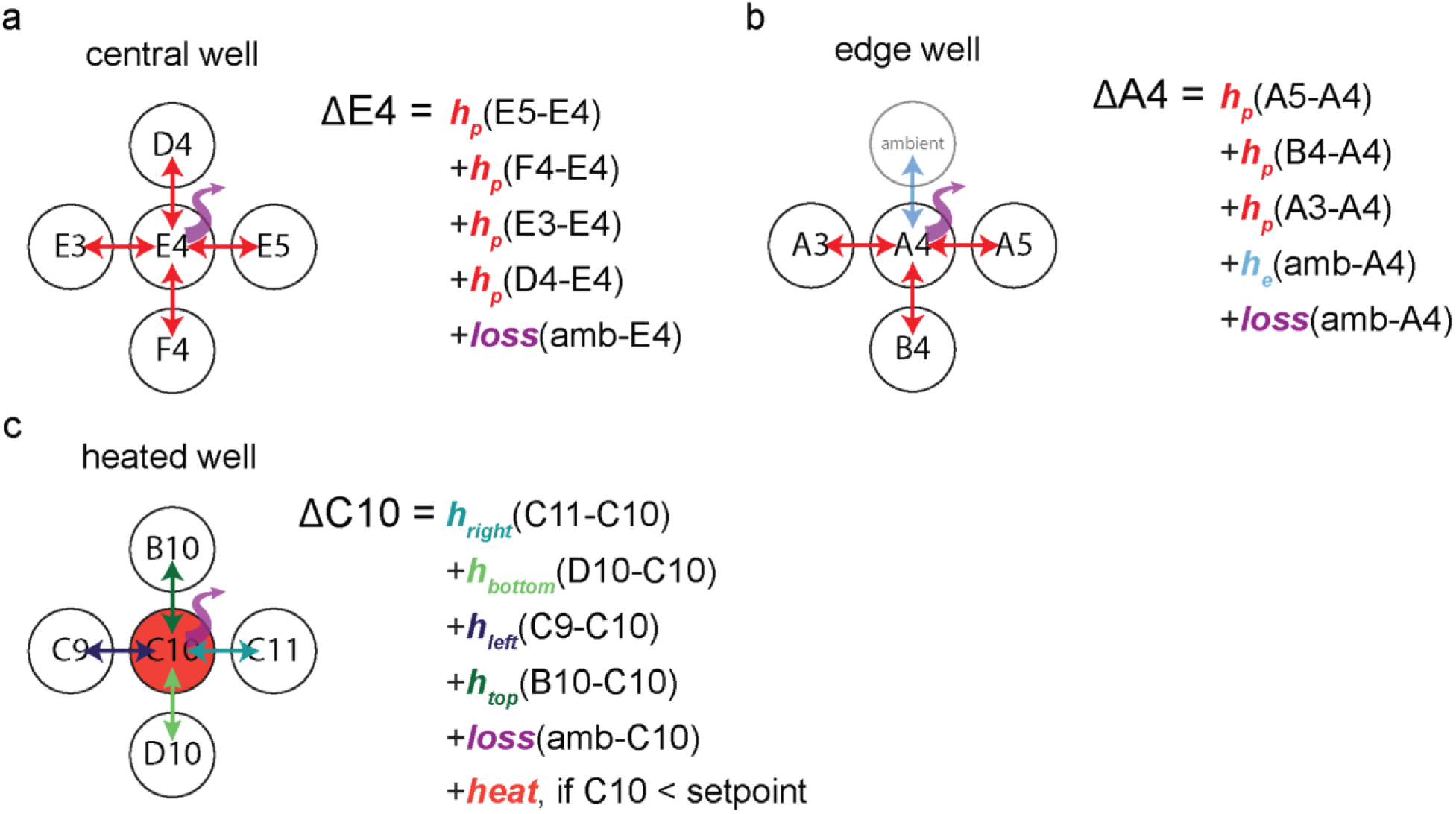
Diagram of all parameters used for model fitting. To fit heat spread across a 96 well plate during thermoPlate operation, wells were categorized into 1 of 3 categories: a) a central well that transfers heat only to neighbor wells and vertically to the environment, b) edge wells that transfer heat to neighbors, vertically to the environment, and horizontally to the environment, and c) heated wells that can transfer heat in any of the ways shown in (a, b) but also receive heat deposition from a thermoPlate heater. For all scenarios, the temperature difference between the well of interest and its neighbors/environment are weighted by fitted parameters to calculate heat transfer.

