## Supplementary material for "Multiplexed dynamic control of temperature to probe and observe mammalian cells": Repository including usage manuals: Assembly Manual.docx

Directions:

1. ***Top Side Placement of Components (Steps 1-10)***. Secure the thermoPlate PCB to a flat surface using tape or L-brackets. Ensure the orientation of the PCB has the top face (with pads) upwards.
2. Secure a solder mask flush against the PCB, taping the edges down to minimize movement. Carefully align the holes to the SMD pads, taking extra care with the smaller pads.
3. Using solder paste (138 °C), deposit a generous amount of solder paste onto one edge of the mask components.
4. Use a solder paste spreader, drag or spread the paste over the mask, pushing down with pressure, creating a thin layer of solder paste to cover all the holes.
5. Remove excess solder paste from the mask using the spreader at a vertical angle.
6. Remove the solder mask carefully, making sure not to smear the paste on the PCB.
7. With a tweezer, place the components onto the appropriate SMD pads on the top side of the thermoplate. With the exception of the multiplexers and shift registers, all components either have no polarity or only align with the pad in one orientation. Shift registers and multiplexers should be placed such that the circular polarity marker is in the top left corner. Do not place any components in the 4 through hole pins in each well position, the barrel jack through holes, or the arduino connector through holes. These will be soldered by hand later. For a diagram of component placement, see the image below:

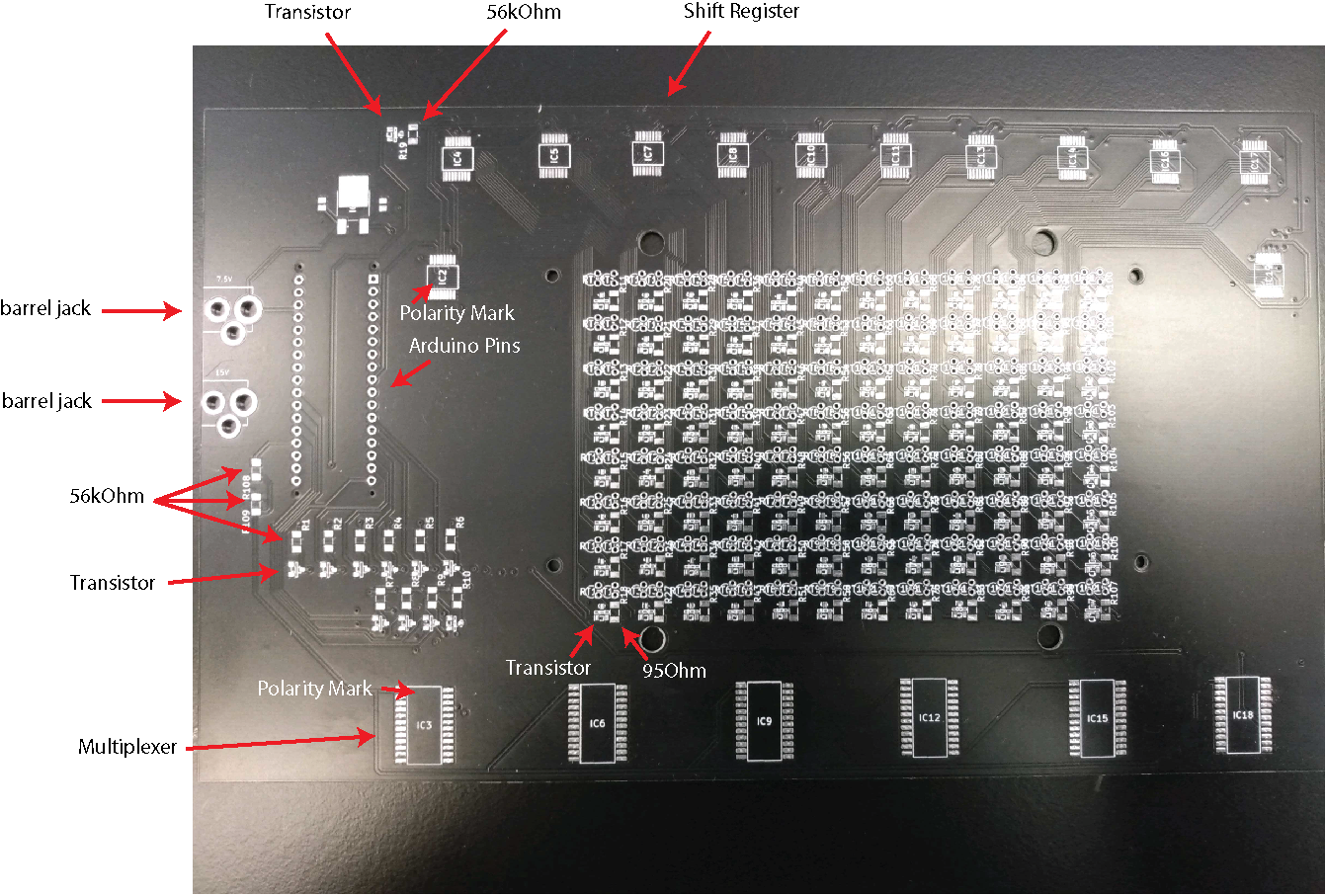

8. Once components are placed, place the PCB on a toaster oven rack.
9. Bake in the toaster oven at 150°C until the solder paste begins to glisten as a shiny silver, indicating its melting. At this point, the components will solder. Use a flashlight to better observe the solder. The larger pads of solder paste may take longer than the smaller ones.
10. Remove the PCB from the toaster oven with heat-resistant gloves and allow it to cool completely on the bench.
11. ***Through Hole Soldering (Steps 11-16).*** With the top face up, place the barrel jack terminals into the PCB with the pins going down. Flip the PCB and solder the pins to the board using a soldering iron.
12. With the bottom face up, insert one pin header into the PCB. Flip the PCB so the pins are pointed up. The black plastic component of the header should be on the side of the PCB with no surface mounted components.
13. Solder one pin to the board with the soldering iron, ensuring the pin header is perpendicular to the PCB. Allow the solder to solidify. The position can be adjusted by holding the iron to the solder to liquify it again.
14. Solder the remaining header pins.
15. Repeat steps 29-30 with the second pin header.
16. Socket pins are permanently soldered into the small through hole pads. These sockets allow the thermistors to be easily inserted onto the board and replaced as needed. Insert the socket pins into their respective through holes such that a single pin is aligned vertically. Once all pins are placed, flip the board, ensuring the pins do not fall out, and place the board upside down on a flat surface, pushing the board against the surface to ensure the socket pins are relatively straight and inserted into their respective through holes. Solder each pin to its pad.
17. ***Thermistor placement and waterproofing (Steps 17-22).*** Insert each thermistor into its appropriate socket (680 Ohm on the left and 330 kOhm on the right).
18. Cut a piece of heat shrink tubing such that when inserted onto the 680 thermistor, it extends just above the center of mass of the thermistor head, but does not extend past the edge of the thermistor. Cut 96 pieces of equal length, testing to ensure the length of each piece is satisfactory as each is cut. Repeat the same steps for the 330 kOhm thermistor.
19. Once all thermistors have been wrapped in heat shrink coating, place the entire assembly into a toaster oven and bake at ~100°C until the heat shrink is tight around the entire thermistor including the pins.
20. Remove the assembly and allow to cool.
21. Spray all thermistors with a generous amount of silicone spray in a fume hood. Move the spray over the entire thermistor area of the board at a steady pace. To apply the appropriate amount of spray, move over the thermistors while spraying at a pace that will cover the entire thermistor area of the board in ~6 seconds of steady movement. Allow to set for 10 minutes. Repeat this process 2-3 times from different angles until all thermistors appear fully and evenly coated. If there is a noticeable buildup of silicone at the base of a thermistor or several thermistors, that area of the board likely does not require further coating even if it has not received a full 3-4 coats.
22. Allow the thermoplate to sit at room temperature for 24 hours or bake at 60°C for 20 minutes to allow silicone curing. Even if the board is cured at 60°C, it is recommended that it is left in a fume hood for 24 hours for the coating fumes to dissipate. It is also recommended to use the User Manual wash protocol before the first use of the thermoPlate to remove any uncured silicone spray.
23. ***3D Printing (Steps 23-24).*** Print the adapter found in the thermoPlate repository and place it such that the thermistors are extending through each of the 96 holes. It is highly recommended to insert the thermistors gently, trying to avoid chipping the waterproof coating. The mounting holes in the adapter should align with the mounting holes of the thermoPlate.
24. Use the screws and bolts found in the parts list to attach the adapter to the thermoPlate.
25. ***Calibration (Steps 25-31).*** Place the thermoPlate on a 96 well plate containing150 µL of PBS in each well and ~100 µL between each well.
26. Place the assembly in a temperature adjustable incubator set to ~25°C and equilibrate overnight (equilibration may be faster if the incubator is already set to a similar temperature).
27. Place two digital thermometers (RC-4 Elitech Digital Temperature Data Logger) inside sealed tubes of water besides the thermoPlate, and cover all components with a plastic box to avoid variation due to airflow.
28. Record the average of the two digital thermometers was recorded as the “true temperature.” Record 5 temperature readings from every well. Average those 5 readings and record that average as the final temperature reading of that well.
29. Set the incubator to increase the ambient temperature to 30°C and allow it to equilibrate for 2 hours. Record readings from the thermoPlate and the digital thermometers in the same manner as above, and repeat this procedure in 5°C increments until 45°C.
30. Using R studio, generate a slope and y-intercept for a line of best fit for the thermoPlate temperature vs true temperature for each well.
31. Import the slopes and y-intercept values into the Arduino file used to control thermoPlate heating (see image below), which is able to automatically incorporate these values into temperature readings. See **Manual** for more details on thermoPlate operation. Slopes should be multiplied by 1000 and Intercepts should be multiplied by -1000 before entering them into the file. Note the orientation of the wells in these arrays:
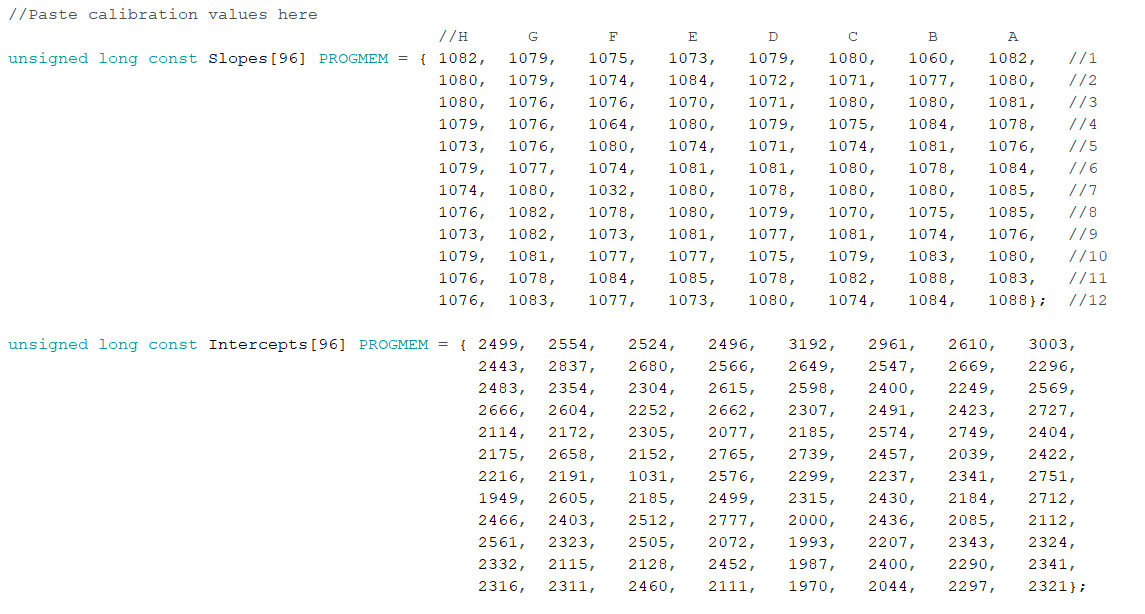
