## Supplementary material for "Multiplexed dynamic control of temperature to probe and observe mammalian cells": Repository including usage manuals: DefaultArduinoScript.ino

```
//time
int t = millis();


//Shift Register pins:
const int latchPin = 5;
const int dataPin = 4;
const int clockPin = 6;
const int Input = A0;
//Multiplexer translation arrays
const int MultiplexerCS[6] = {7, 8, 9, 10, 11, 12};
//S0 should be first
const int MultiplexerInputs[4] = {A1,A2,A3,A4};
//Master Control
const int Master = 13;
double Vin = 12.500;
double Rd = 56000.000;
double B2585 = 4570.000;
double R25 = 330000.000;
double RT = 306000.000;
double R1 = 100000.000;

//PID constants
double kp = 100;
double ki = .0002;
double kd = .4;

//PID variables
unsigned long currentTime;
unsigned long previousTime[96];
double elapsedTime;
double error;
double lastError[96];
double cumError[96]; 
double rateError;
double expectedError;

double DecimalCorrection = 100.000;

//Enter temperatures desired for the first phase below
//3700 corresponds to 37.00 degrees
//3750 corresponds to 37.5 degrees. Etc.

                                            //8  7  6  5  4  3  2  1
uint16_t const ON_SetPoint[96] PROGMEM = {  0, 0, 0, 0, 0, 0, 0, 0, //1
                                            0, 0, 0, 0, 0, 0, 0, 0, //2
                                            0, 0, 0, 0, 0, 0, 0, 0, //3
                                            0, 0, 0, 0, 0, 0, 0, 0, //4
                                            0, 0, 0, 0, 0, 0, 0, 0, //5
                                            0, 0, 0, 0, 0, 0, 0, 0, //6
                                            0, 0, 0, 0, 0, 0, 0, 0, //7
                                            0, 0, 0, 0, 0, 0, 0, 0, //8
                                            0, 0, 0, 0, 0, 0, 0, 0, //9
                                            0, 0, 0, 0, 0, 0, 0, 0, //10
                                            0, 0, 0, 0, 0, 0, 0, 0, //11
                                            0, 0, 0, 0, 0, 0, 0, 0, //12
                                          };

//Enter temperatures desired for the second phase below
uint16_t const OFF_SetPoint[96] PROGMEM =  {0, 0, 0, 0, 0, 0, 0, 0, //1
                                            0, 0, 0, 0, 0, 0, 0, 0, //2
                                            0, 0, 0, 0, 0, 0, 0, 0, //3
                                            0, 0, 0, 0, 0, 0, 0, 0, //4
                                            0, 0, 0, 0, 0, 0, 0, 0, //5
                                            0, 0, 0, 0, 0, 0, 0, 0, //6
                                            0, 0, 0, 0, 0, 0, 0, 0, //7
                                            0, 0, 0, 0, 0, 0, 0, 0, //8
                                            0, 0, 0, 0, 0, 0, 0, 0, //9
                                            0, 0, 0, 0, 0, 0, 0, 0, //10
                                            0, 0, 0, 0, 0, 0, 0, 0, //11
                                            0, 0, 0, 0, 0, 0, 0, 0, //12
                                          };


//Enter the duration of the first phase (minutes)
unsigned long const ON_Time[96] PROGMEM = { 0, 0, 0, 0, 0, 0, 0, 0, //1
                                            0, 0, 0, 0, 0, 0, 0, 0, //2
                                            0, 0, 0, 0, 0, 0, 0, 0, //3
                                            0, 0, 0, 0, 0, 0, 0, 0, //4
                                            0, 0, 0, 0, 0, 0, 0, 0, //5
                                            0, 0, 0, 0, 0, 0, 0, 0, //6
                                            0, 0, 0, 0, 0, 0, 0, 0, //7
                                            0, 0, 0, 0, 0, 0, 0, 0, //8
                                            0, 0, 0, 0, 0, 0, 0, 0, //9
                                            0, 0, 0, 0, 0, 0, 0, 0, //10
                                            0, 0, 0, 0, 0, 0, 0, 0, //11
                                            0, 0, 0, 0, 0, 0, 0, 0, //12

                                          };
                                          
//Enter the duration of the first phase (minutes)
unsigned long const OFF_Time[96] PROGMEM = {0, 0, 0, 0, 0, 0, 0, 0, //1
                                            0, 0, 0, 0, 0, 0, 0, 0, //2
                                            0, 0, 0, 0, 0, 0, 0, 0, //3
                                            0, 0, 0, 0, 0, 0, 0, 0, //4
                                            0, 0, 0, 0, 0, 0, 0, 0, //5
                                            0, 0, 0, 0, 0, 0, 0, 0, //6
                                            0, 0, 0, 0, 0, 0, 0, 0, //7
                                            0, 0, 0, 0, 0, 0, 0, 0, //8
                                            0, 0, 0, 0, 0, 0, 0, 0, //9
                                            0, 0, 0, 0, 0, 0, 0, 0, //10
                                            0, 0, 0, 0, 0, 0, 0, 0, //11
                                            0, 0, 0, 0, 0, 0, 0, 0, //12
                                      };


//Place Slopes of calibration curves below multipled by 1000                          
unsigned long const Slopes[96] PROGMEM = { 1082,  1079,   1075,   1073,   1079,   1080,   1060,   1082,   
                                           1080,  1079,   1074,   1084,   1072,   1071,   1077,   1080,   
                                           1080,  1076,   1076,   1070,   1071,   1080,   1080,   1081,   
                                           1079,  1076,   1064,   1080,   1079,   1075,   1084,   1078,   
                                           1073,  1076,   1080,   1074,   1071,   1074,   1081,   1076,   
                                           1079,  1077,   1074,   1081,   1081,   1080,   1078,   1084,   
                                           1074,  1080,   1032,   1080,   1078,   1080,   1080,   1085,   
                                           1076,  1082,   1078,   1080,   1079,   1070,   1075,   1085,   
                                           1073,  1082,   1073,   1081,   1077,   1081,   1074,   1076,   
                                           1079,  1081,   1077,   1077,   1075,   1079,   1083,   1080,   
                                           1076,  1078,   1084,   1085,   1078,   1082,   1088,   1083,   
                                           1076,  1083,   1077,   1073,   1080,   1074,   1084,   1088};

//Place the y-intercepts of the calibration curves below
//Mulitplied by -1000
unsigned long const Intercepts[96] PROGMEM = { 2499,  2554,   2524,   2496,   3192,   2961,   2610,   3003,   
                                               2443,  2837,   2680,   2566,   2649,   2547,   2669,   2296,  
                                               2483,  2354,   2304,   2615,   2598,   2400,   2249,   2569,   
                                               2666,  2604,   2252,   2662,   2307,   2491,   2423,   2727,  
                                               2114,  2172,   2305,   2077,   2185,   2574,   2749,   2404,   
                                               2175,  2658,   2152,   2765,   2739,   2457,   2039,   2422,   
                                               2216,  2191,   1031,   2576,   2299,   2237,   2341,   2751,   
                                               1949,  2605,   2185,   2499,   2315,   2430,   2184,   2712,   
                                               2466,  2403,   2512,   2777,   2000,   2436,   2085,   2112,   
                                               2561,  2323,   2505,   2072,   1993,   2207,   2343,   2324,   
                                               2332,  2115,   2128,   2452,   1987,   2400,   2290,   2341,   
                                               2316,  2311,   2460,   2111,   1970,   2044,   2297,   2321};

//const word Zero = 0;
byte ShiftChan[12] = {B00000000, B00000000, B00000000, B00000000, B00000000, B00000000, B00000000, B00000000, B00000000, B00000000, B00000000, B00000000};


void setup() {
  // put your setup code here, to run once:
  pinMode(latchPin, OUTPUT);
  pinMode(clockPin, OUTPUT);
  pinMode(dataPin, OUTPUT);
  pinMode(7, OUTPUT);
  pinMode(8, OUTPUT);
  pinMode(9, OUTPUT);
  pinMode(10, OUTPUT);
  pinMode(11, OUTPUT);
  pinMode(12, OUTPUT);
  pinMode(A1, OUTPUT);
  pinMode(A2, OUTPUT);
  pinMode(A3, OUTPUT);
  pinMode(A4, OUTPUT);
  pinMode(Master,OUTPUT);
  pinMode(Input, INPUT);
  Serial.begin(9600);
  
  //Turn off all the multiplexers now so that we can turn them on one at a time later.
  for (int i = 0; i < 6; i++){
    digitalWrite(MultiplexerCS[i],LOW);
  }
  updateRegisters();
  digitalWrite(Master, HIGH);
  delay(2000);
}

unsigned long CurrentTime;
unsigned long CurrentTimeS;
unsigned long CurrentTimeM;
unsigned long LastTime = 0;

double CurrentTemp;
uint8_t Duty;
uint8_t Duties[96];
double Current_SetPoint;
unsigned long Current_ON;
unsigned long Current_OFF;

//Inter determines the number of PID loops performed
//before temperatures are printed. Each loop takes
//~2 seconds, meaning an Inter = 15 produces readings
//every 30s
int Inter = 15;
int tracker = Inter;
int MinTemp = 5.00;

double slope;
double intercept;

void loop() {
  CurrentTime=millis();
  CurrentTimeS = CurrentTime/1000;
  CurrentTimeM = CurrentTimeS/60;
  if (CurrentTime>LastTime+1999|LastTime==0) {
      for (int i = 0; i<96; i++) {
        slope = (pgm_read_word_near(Slopes+i))/1000.0000;
        intercept = (pgm_read_word_near(Intercepts+i))/1000.0000;
        CurrentTemp = CalcTemp(ReadPin(i))*slope-intercept;
        
        if (CurrentTemp<MinTemp) {
          CurrentTemp=MinTemp;
        }
        Current_ON = pgm_read_word_near(ON_Time+i);
        Current_OFF = pgm_read_word_near(OFF_Time+i);
        if (Current_ON+Current_OFF == 0) {
          Current_OFF = 1;
        }
        if (CurrentTimeM%(Current_ON+Current_OFF) < Current_ON){
          Current_SetPoint = (pgm_read_word_near(ON_SetPoint+i))/DecimalCorrection;
        }
        else {
          Current_SetPoint = (pgm_read_word_near(OFF_SetPoint+i))/DecimalCorrection;
        }
        Duty = computePID(CurrentTemp, Current_SetPoint, i);
        Duties[i] = Duty;
        if (tracker==Inter) {
           Serial.print(CurrentTemp);
           Serial.print(", ");
        }
      }
      if (tracker==Inter) {
         Serial.print(millis());
         Serial.print(", ");
         Serial.print((RT/R1)*5.000*analogRead(A5)/1023.000);
         Serial.print(", ");
         Serial.print(B2585);
         Serial.print(", ");
         Serial.print(CurrentTimeM);
         Serial.println();
         tracker=0;
      }
      LastTime=millis();
      tracker+=1;
  }
  for (int i=0; i<96; i++) {
    if (CurrentTime%100<Duties[i]) {
      updateBits(i, 1);
    }
    else {
      updateBits(i, 0);
    }
  }
}


void updateRegisters(){
  digitalWrite(latchPin, LOW);
  for (int i=11; i>-1; i--) {
    shiftOut(dataPin, clockPin, MSBFIRST, ShiftChan[i]);
  }
  digitalWrite(latchPin, HIGH);
}


void updateBits(int Well, int OnOff) {
  int ListIndex = Well/8;
  int BitIndex = Well%8;
  byte Output = ShiftChan[ListIndex];
  if (OnOff == 0) {
    bitClear(Output, BitIndex);
  }
  else {
    bitSet(Output, BitIndex);
  }
  ShiftChan[ListIndex] = Output;
  updateRegisters();
}


double ReadPin(int Well){
    int CSIndex = Well/16;
    int Pin = Well%16;
    for (int i=0; i<6; i++) {
      digitalWrite(MultiplexerCS[i], LOW);
    }
    
    for (int i = 0; i<4; i++){
      byte state = bitRead(Pin, i);
      digitalWrite(MultiplexerInputs[i], !state);
    }
    digitalWrite(MultiplexerCS[CSIndex], HIGH);
    delay(1);
    double Reading = analogRead(Input);
    digitalWrite(MultiplexerCS[CSIndex], LOW);
    for (int i = 0; i<4; i++){
      digitalWrite(MultiplexerInputs[i], 0);
    }
    //Reading = Reading-CalSubtract[Well];
    return Reading;
}


double CalcTemp(double Reading) {
  double Vo = 5.000*Reading/1023.000;
  double ActualVin = (RT/R1)*5.000*analogRead(A5)/1023.000;
  double R2 = ActualVin/Vo*Rd-Rd;
  double T2 = (1.000/(-1.000*(log(R25/R2))/B2585 + 1.000/(273.000+25.000)))-273.000;
  return T2;
}

 
double computePID(double inp, double SetPoint, int i){     
        currentTime = millis();                //get current time
        elapsedTime = (double)(currentTime - previousTime[i]);        //compute time elapsed from previous computation
        error = SetPoint - inp;                                // determine error
        cumError[i] = cumError[i]+ (error * elapsedTime);                // compute integral
        if (cumError[i]*ki > 100) {
          cumError[i] = 100/ki;
        }
        if (cumError[i]*ki < -100) {
          cumError[i] = -100/ki;
        }
        rateError = (error - lastError[i])/elapsedTime;   // compute derivative
        expectedError = error+error-lastError[i];
        double out = kp*error + ki*cumError[i] + kd*rateError; //+ke*expectedError;                //PID output               
        lastError[i] = error;                                //remember current error
        previousTime[i] = currentTime;                        //remember current time
        if (out<-100) {
          out = -100.000;
        }
        if (out>100) {
          out = 100.000;
        }
        out = (out/2.000)+50.000;
        return out;                                        //have function return the PID output
}
```
