## Supplementary material for "Multiplexed dynamic control of temperature to probe and observe mammalian cells": Repository including usage manuals: User Manual.docx

**Usage Protocol:**

1. Download the Arduino IDE software: <https://www.arduino.cc/en/software>. (Version 1.8.19 is preferred.)
2. Upon downloading, install this program and open the file “Default Arduino Script.ino” found in the repository. Ensure the USB cable (found in parts list) can reach between the computer being used and the location where the thermoPlate will be placed.
3. Prior to mounting and using the thermoplate, first program the desired protocol. Locate the two arrays named ON_SetPoint and set each well to the desired temperature multiplied by 100 (37.5C is input as 3750). These will be the set temperature of the first temperature phase
4. Locate the OFF_SetPoint array and input temperatures in the same format. This will be the temperature setpoint of the second phase. The thermoPlate cycles between each temperature phase indefinitely.

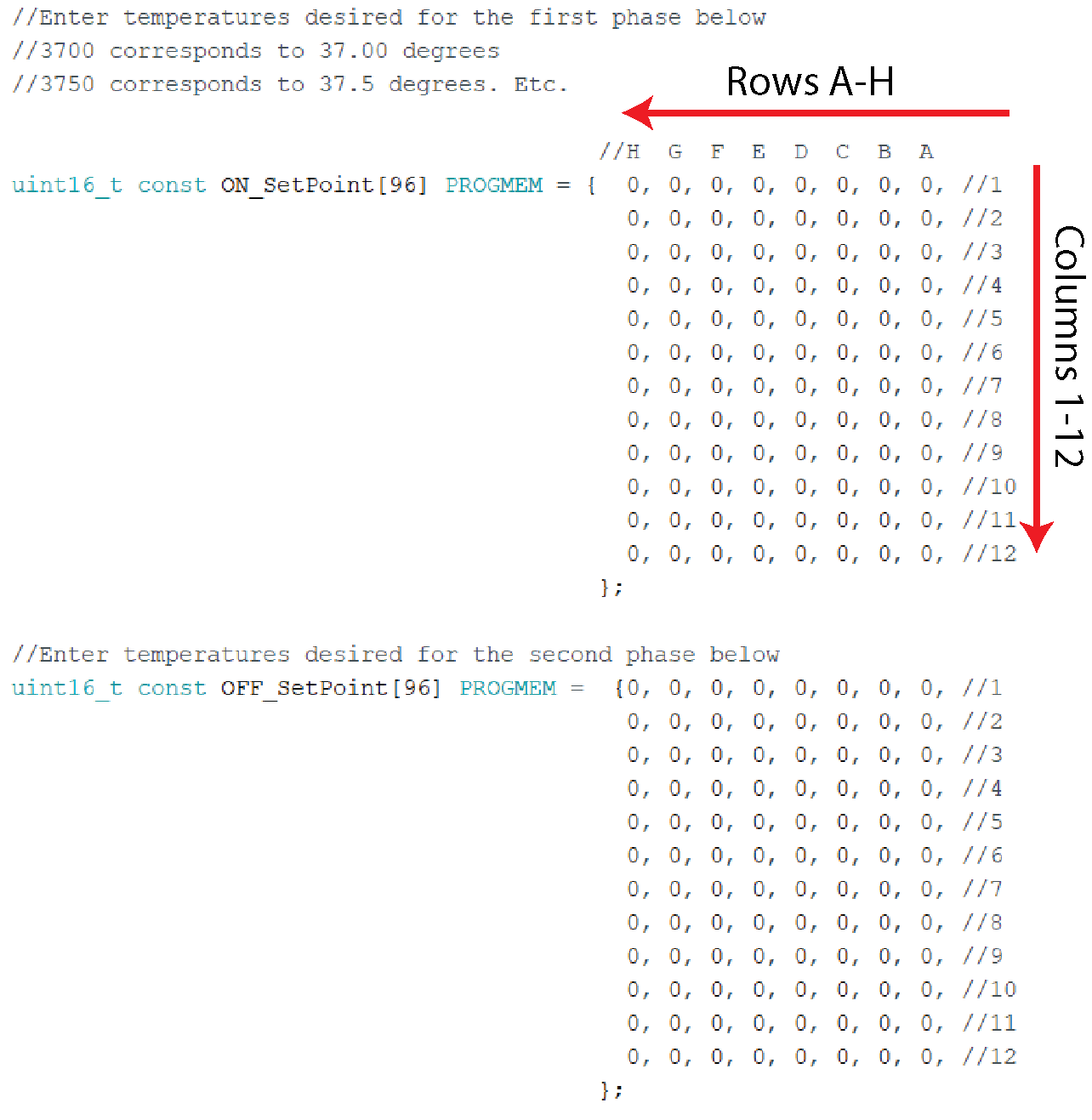

5. Locate the array ON_Time and OFF_Time. Input the time, in minutes, that each phase will last before transitioning to the next phase.
6. Locate the line below the comment “Master Heating Switch”. Set this to 0 as seen below:
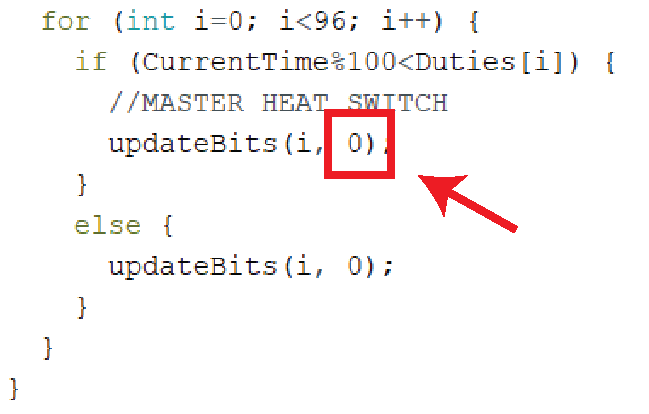

Setting this number to 0 places the thermoPlate in “measurement mode.” If only measurements and no heating is desired (for example, during calibration), maintain a 0 in this position for the rest of the protocol. However, if heating is desired, it is still recommended to start the arduino in measurement mode. Otherwise, the thermoPlate will run the last protocol that was uploaded immediately upon plugging in the 12V power cable.

7. Sterilize the thermoPlate by spraying generously with ethanol and blow drying with pressurized air. Do not allow the ethanol to remain on the device for >5 minutes, as this may corrode the waterproof coating.
8. Insert the thermoPlate into a 96 well plate. The plate should have ~150 µL of fluid in each well and ~100 µL of PBS between each well
9. Plug in the 7.5V power adapter to the 7.5V barrel jack.
10. Plug in the USB cable to the the Arduino and to a computer
11. Upload the Arduino script to the Arduino (ensure that the correct USB port is selected in the Arduino software under Tools->Port and ensure that the Board is set to Arduino Micro under Tools->Board.
12. Open the Arduino Serial Monitor under Tools->Serial monitor. Ensure that temperatures are printing to the Serial Monitor (they will read 5.00 until the final power cord is plugged in.
13. Plug in the final 12V power adapter to the 12V barrel jack. Readings should now read the ambient room temperature
14. When the protocol is ready to be run, change the Master Heat Switch from 0 to 1 and reupload. The protocol will immediately start.

NOTE: It is recommended to always change the Master Heat Switch to 0 and upload to the Arduino before plugging in the 12V plug. This will prevent the thermoPlate from heating wells using a previously uploaded protocol.

**Washing Protocol:**

1. When the protocol is complete, remove the thermoPlate from the culture plate. Some thermistors may have shifted during the protocol, making it difficult to remove. In this case, gently prise the thermoPlate off the plate along the corners and edges until it begins to come away from the plate.
2. Fill a micropipette tip box (or similar sized vessel) with DI water. Place a magnetic stir bar in the water, and place the vessel on a stirring platform.
3. Insert the thermoPlate into the water such that each thermistor is completely submerged.
4. Turn on the stirring function to remove media and other contaminants from the thermistors
5. Wash for ~10 min - 2 hrs and blow dry using pressurized air from an air spout. UV sterilization is also recommended after each use.
