## Supplementary material for "Multiplexed dynamic control of temperature to probe and observe mammalian cells": Files for running the thermoPlate GUI: app.R

```
#
# This is a Shiny web application. You can run the application by clicking
# the 'Run App' button above.
#
# Find out more about building applications with Shiny here:
#
#    http://shiny.rstudio.com/
#
# #install.packages("matlab")
# #install.packages("openxlsx")
# install.packages("tidyverse")
# install.packages("gganimate")
# install.packages("gifski")
# install.packages("vroom")
# install.packages("png")
library(openxlsx)
library(matlab)
library(shiny)
library(ggplot2)
library(dplyr)
library(tidyverse)
library(gganimate)
library(gifski)
library(vroom)
library(png)

source("SampleInputFile.R")
source("ReadInputFile.R")
source("ArduinoToApp.R")
source("ArduinoToModel.R")
source("PlotInputMaker.R")
source("ModelToApp.R")
source("RunModelV2.R")
source("wellPlatePlot.R")
source("wellPlateAnim.R")
source("heatErrorCheck.R")


# Define UI for app that runs simulation ----
ui <- fluidPage(
    titlePanel("Thermoplate Heating GUI"),
    
    sidebarLayout(
        sidebarPanel(position = "left",
            h5("1. Download .xlsx template to specify experimental wells, times, and temps"),
            
            downloadLink('downloadData', 'Download input template (.xlsx)'),
            br(),
            br(),

            fileInput("upload", h5("2. Upload xlsx with wells, times, and temps"), accept = ".xlsx"),

            numericInput("length",
                         h5("3. Input length of experiment (mins)"),
                         value = 30,
                         min = 0,
                         max = 1000),
            br(),
            
            h5("4. When ready to run simulation, press Simulate"),
            actionButton("start",
                         "Simulate")
        ),
        
        mainPanel(
                  h4("Planned heating profile"),
                  plotOutput("plot"),
                  uiOutput("slider"), #placeholder for slider once upload data
                  
                  h4("Heating simulation"),
                  plotOutput("animation"),
                  )
    )
)


# Define server logic required to run a simulation ----
server <- function(input, output) {
  timestep = 25 # time resolution (seconds) that will be plotted in output. i.e. each frame corresponds to this many seconds.
  
  #make a lookuptable that matches rows, cols, and WellID
  ColNum = c(rep(1,8),rep(2,8),rep(3,8),rep(4,8),rep(5,8),rep(6,8),rep(7,8),rep(8,8),rep(9,8),rep(10,8),rep(11,8),rep(12,8))
  RowNum = rep(1:8,12)
  wellLUT = data.frame(ColNum, RowNum) %>% 
    mutate(RowName = 
             case_when(RowNum == 1 ~"A",
                       RowNum == 2 ~"B",
                       RowNum == 3 ~"C",
                       RowNum == 4 ~"D",
                       RowNum == 5 ~"E",
                       RowNum == 6 ~"F",
                       RowNum == 7 ~"G",
                       RowNum == 8 ~"H")) %>% 
    mutate(WellID = paste(RowName, ColNum, sep = ""))
  rownames(wellLUT) <- NULL
  rm(ColNum,RowNum)
    
    output$downloadData <- downloadHandler(
          filename = 'SampleInput.xlsx',
          content = function(con){
            SampleInputFile(con)
          }
        )
    
    #store experimental length
    ExperimentLength = reactive({
      input$length
    })
    
    #import well, time, temp .csv
    UserInput = reactive({
      ReadInputFile(input$upload$datapath)
    })
    
    # Reactive expression to generate a unique key for each upload
    uploadKey <- reactive({
      if (is.null(input$upload)) {
        return(NULL)
      } else {
        # Using file modification time
        file.info(input$upload$datapath)$mtime
      }
    })
    
    # Convert User Input to that required by App
    AppInput = reactive({
      if(!is.null(UserInput())){
        ArduinoToApp(wellLUT,ExperimentLength(),UserInput())
      }
    })
    
    # Convert User Input to that required by App's plots
    WellPlotInput = reactive({
      if(!is.null(AppInput())){
        PlotInputMaker(wellLUT,ExperimentLength(),AppInput())
      }
    })
    
    # Convert User Input to that required by simulation
    ModelInput = reactive({
      if(!is.null(UserInput())){
        ArduinoToModel(ExperimentLength(),timestep,UserInput())
      }
    })
    
    
    #show slider once data is input
    output$slider <- renderUI({
        req(input$upload)
        sliderInput("times", 
                    label = "Run time",
                    min = 0, 
                    max = ExperimentLength(), 
                    value = 0,
                    step = 1,
                    animate = TRUE)
    })
    
    
    #display input table
    output$upload_table <- renderTable(
        head(AppInput())
    )
    
    #create interactive plate with heats and numbers
    wellPlot = reactive({
        if(!is.null(input$upload)){
            wellPlatePlot(WellPlotInput(), input$times,ExperimentLength())#generate a plot
        }
      })
    
    #display the plot
    output$plot = renderCachedPlot({
        wellPlot()
       },
        sizePolicy = sizeGrowthRatio(width = 600, 
                                     height = 400, 
                                     growthRate = 1),
        # height = 400,
        # width = 600,
        cacheKeyExpr = list(input$times, uploadKey())
    )
    
    # Run the simulaiton
    heatSimulation = eventReactive(input$start,{
        # Create a Progress object
        progressModel <- shiny::Progress$new()
        progressModel$set(message = "Computing", value = 0)
        # Close the progress when this reactive exits (even if there's an error)
        on.exit(progressModel$close())
        
        # Create a callback function to update progress.
        # Each time this is called:
        # - If `value` is NULL, it will move the progress bar 1/5 of the remaining
        #   distance. If non-NULL, it will set the progress to that value.
        # - It also accepts optional detail text.
        updateProgress <- function(value = NULL, detail = NULL) {
            if (is.null(value)) {
                value <- progressModel$getValue()
            }
            progressModel$set(value = value, detail = detail)
        }
        
        heatMod = heating_eval(ModelInput(), timestep, updateProgress) 
        heatMod = ModelToApp(wellLUT,heatMod,timestep,ExperimentLength())
        # Create a Progress object for animation
        progressAnim <- shiny::Progress$new()
        progressAnim$set(message = "Animating...", value = NULL)
        # Close the progress when this reactive exits (even if there's an error)
        on.exit(progressAnim$close(), add = TRUE)
        
        #check if some wells are hotter than programmed by user
        heatMod_wErr = heatErrorCheck(heatMod, WellPlotInput(),ExperimentLength()) 
        wellPlateAnim(heatMod_wErr)
    })
        
    
    #output the sim gif
    output$animation = renderImage({
        #specify which image to use (placeholder image if no data yet)
        if(is.null(input$upload) | input$start == 0){
            list(src = "simPlaceHolder.png",  #call the reactive gif file here
                 contentType = 'image/png',
                 width = 600,
                 height = 400,
                 alt = "This is alternate text")
          }else{list(src = heatSimulation(),  #call the reactive gif file here
                                     contentType = 'image/gif',
                                     width = 600,
                                     height = 400,
                                     alt = "This is alternate text")
              }
    },deleteFile = ifelse(is.null(input$upload),FALSE,TRUE)
)
}

# Run the application 
shinyApp(ui = ui, server = server)
```
