## Supplementary material for "Multiplexed dynamic control of temperature to probe and observe mammalian cells": Files for running the thermoPlate GUI: ArduinoToApp.R

```
ArduinoToApp <- function(wellLUT, ExperimentLength, Inputs){
  # This function converts the user input to one usable by the app.
  
  #Inputs will be a list of arrays with names OnTime, OffTime, OffTemp, OnTemp.
  for (i in 1:length(Inputs)){
    x = names(Inputs)[i]
    assign(x, Inputs[[i]], pos = -1)
  }

  OnWells = (OnTemp == 0) + (OffTemp == 0) <= 1
  OnWellIndices = which(OnWells)
  inputDF = data.frame(matrix(ncol = 7, nrow = 0))
  for (w in OnWellIndices){
    cLen = OnTime[w] + OffTime[w]
    if (cLen == 0){
      stop(paste("Well", wellLUT$WellID[w], "is set so that both OnTime and OffTime are 0. This is not possible.", sep = " "))
    }
    sTimes = seq(0,ExperimentLength, by=cLen)
    sTimes = append(sTimes,sTimes + OnTime[w])
    sTimes = sort(sTimes)
    sTimes = sTimes[sTimes < ExperimentLength]
    eTimes = append(sTimes[2:length(sTimes)],ExperimentLength)
    Temperatures = rep(c(OnTemp[w],OffTemp[w]),floor(length(sTimes)/2))
    if (length(Temperatures) < length(sTimes)){
      Temperatures = append(Temperatures,OnTemp[w])
    }
    temp = suppressWarnings(cbind(wellLUT[w,],sTimes,eTimes,Temperatures))
    inputDF = rbind(inputDF,temp)
  }
  colnames(inputDF) <- c('ColNum','RowNum','RowName','WellID','start','stop','temp')
  return(inputDF)
}
```
