## Supplementary material for "Multiplexed dynamic control of temperature to probe and observe mammalian cells": Files for running the thermoPlate GUI: ArduinoToModel.R

```
ArduinoToModel <- function(ExperimentLength, timestep, Inputs){
  # This function converts the user input to one usable by the simulation.
  
  ExperimentLength = ExperimentLength*60
  #Inputs will be a list of arrays with names OnTime, OffTime, OffTemp, OnTemp.
  for (i in 1:length(Inputs)){
    x = names(Inputs)[i]
    assign(x, Inputs[[i]], pos = -1)
  }
  OnTime <- OnTime*60
  OffTime <- OffTime*60
  
  t = seq(0,ExperimentLength+timestep, by=timestep)
  
  # Create a 3D array to store temperature values
  TargetTemps <- array(NA, dim=c(8, 12, length(t)))
  
  # Fill the 3D array with temperature values
  for (i in 1:length(t)){
    denominator = (OnTime + OffTime)
    denominator[denominator == 0]=1
    WellIsOn = t[i]%%(denominator) < OnTime
    TargetTemps[,,i] = OnTemp*WellIsOn + OffTemp*(!WellIsOn)
  }
  return(TargetTemps)
}
```
