## Supplementary material for "Multiplexed dynamic control of temperature to probe and observe mammalian cells": Files for running the thermoPlate GUI: heatErrorCheck.R

```
heatErrorCheck = function(heatMod, WellPlotInput,ExperimentLength){
  # Flags errors for making the simulation
  
  tol = 0.5 #error tolerance in degrees Celsius
  
  Inputs <- WellPlotInput %>%
    arrange(WellID,start) %>%
    group_by(WellID) %>%
    mutate(
      CoolingPhase = temp-lag(temp) < 0,
      CoolingPhase = coalesce(CoolingPhase,FALSE),
      CoolingPhase = if_else(start==0 & stop==ExperimentLength & temp==0,NA,CoolingPhase)
    ) %>%
    ungroup() %>%
    select(WellID,start,stop,Target=temp,CoolingPhase)
  
  
  result <- heatMod %>%
    inner_join(Inputs, by = "WellID", relationship = "many-to-many") %>%
    filter(time >= 60*start, time < 60*stop) %>%
    arrange(WellID,time) %>%
    group_by(WellID) %>%
    mutate(
      # dTdt = (lead(temp)-temp)/(lead(time)-time),
      # dTdt = (temp-lag(temp))/(time-lag(time)),
      # dTdt = coalesce(dTdt,0),
      # error = case_when(
      ## Detecting error when the well is heating and over its setpoint
      #   !CoolingPhase & (temp - Target > tol) ~ 1,
      ## Detecting error when the well is in a cooling phase, but not cooling fast enough (by linear extrapolation) to reach its setpoint by the end of the phase.
      #   CoolingPhase & (temp - Target > tol) & (temp-Target + dTdt*(60*stop-time)>tol) ~ 1,
      ## Detecting error if well is in a cooling phase, but is actually rising in temperature above setpoint
      #   # CoolingPhase & (temp - Target > tol) & (dTdt > 0) ~ 1,
      #   .default = 0
      # )
      
      ## Detecting error anytime a well is over its setpoint
      error = case_when(
        !is.na(CoolingPhase) & (temp - Target > tol) ~1,
        .default = 0
      )
    ) %>%
    ungroup() %>%
    select(WellID, ave, time, RowName, RowNum, ColNum, temp, error)

  return(result)
}
```
