## Supplementary material for "Multiplexed dynamic control of temperature to probe and observe mammalian cells": Files for running the thermoPlate GUI: ModelToApp.R

```
ModelToApp <- function(template,timecourse,timestep,ExperimentLength){
  # Converts the simulation output to one that can be plotted by the app
  ExperimentLength = ExperimentLength*60
  t = seq(0,ExperimentLength+timestep, by=timestep)
  output <- data.frame(matrix(ncol = 4, nrow = 0))
  for (i in 1:length(t)){
    time = rep(t[i],96)
    temp = timecourse[,,i]
    temp = reshape(temp,c(96,1))
    ave = temp
    Entry = cbind(template,time,temp,ave)
    output = rbind(output,Entry)
  }
  output <- output[, c('WellID','ave','time','RowName','RowNum','ColNum','temp')]
  return(output)
}
```
