## Supplementary material for "Multiplexed dynamic control of temperature to probe and observe mammalian cells": Files for running the thermoPlate GUI: PlotInputMaker.R

```
PlotInputMaker = function(wellLUT,ExperimentLength,AppInput){
  # Makes the planned heat profile preview's input
  WellsNotUsed <- wellLUT %>% filter(!(WellID %in% AppInput$WellID))
  start <- rep(0,length(WellsNotUsed$ColNum))
  stop <- rep(ExperimentLength,length(WellsNotUsed$ColNum))
  temp <- rep(0,length(WellsNotUsed$ColNum))
  WellPlotInput <- cbind(WellsNotUsed,start,stop,temp)
  WellPlotInput <- rbind(AppInput,WellPlotInput)
  return(WellPlotInput)
}
```
