## Supplementary material for "Multiplexed dynamic control of temperature to probe and observe mammalian cells": Files for running the thermoPlate GUI: ReadInputFile.R

```
ReadInputFile <- function(filename){
    # Load the workbook
    wb <- loadWorkbook(filename)
    
    # Get the names of all worksheets
    sheet_names <- names(wb)
    
    # Create a list to hold arrays for each worksheet
    arrays_list <- list()
    
    # Iterate over each worksheet and read its content
    for(sheet in sheet_names) {
      data <- read.xlsx(filename, sheet = sheet, colNames = FALSE)
      arrays_list[[sheet]] <- array(unlist(data), dim = c(8,12))
    }
    
    return(arrays_list)
}
```
