## Supplementary material for "Multiplexed dynamic control of temperature to probe and observe mammalian cells": Files for running the thermoPlate GUI: README.txt

Purpose of the App:
This app allows users of the thermoPlate device to run simulations of their planned heating timecourses in order to see if the device can feasibly maintain the desired temperature differentials between wells. 

Initial Launch Instructions:
1) Extract all files into a single folder.
2) Ensure that R is properly installed and updated. (This app was made on a computer using R 4.3.0).
3) Open the file named "app.R" in RStudio.
4) Ensure that all necessary packages are installed. This can be done by uncommenting and running the "install.packages" commands at the top of the "app.R" file. After running once, these lines can be commented once more and need not be run again.
5) If some or all packages are already installed, ensure they are updated

Running a Simulation:
1) After completing the steps above, click on the "Run App" button on the top right of the panel displaying the R script. The app window will display.
2) Upload a properly formatted input file. A sample input file can be obtained by clicking "Download input template .xlsx" in the app window. The format is described in greater detail in the section below.
3) Input the length of the experiment in minutes.
4) [Optional] Use the "Run time" slider to preview the planned heating profile.
5) Click Simulate.
6) Once complete, the output of the simulation will display as a GIF in the "Heating simulation" panel. Wells whose temperature exceed their setpoint will be circled in red for as long as their temperature exceeds their setpoint.

Input Format:
The app takes its input in the form of a .xlsx file with 4 worksheets titled "OnTemp", "OffTemp", "OnTime", and "OffTime". All worksheets hold data in 8 rows and 12 columns, where cell A1 corresponds to Well A1 of a 96 well plate and cell L1 corresponds to Well A12. In the "OnTemp" and "OffTemp" worksheets, please enter temperatures in Degrees Celsius above the intended Ambient Temperature. In the "OnTime" and "OffTime" worksheets, please enter the amount of time (in minutes) you want the the well to stay in the On and Off phases, respectively. Wells enter the On phase first, and cycle between the On and Off phases for the duration of the simulation.
Please note that the "Planned Heating Profile" plot will only display wells whose temperatures are not set to 0 over all time. Similarly, the simulation only flags errors in wells whose temperatures are not set to 0 over all time. 
