## Supplementary material for "Multiplexed dynamic control of temperature to probe and observe mammalian cells": Files for running the thermoPlate GUI: RunModelV2.R

```
heating_eval <- function(on_wells, timestep, updateProgress = NA) {
  # Runs the simulation
  parameters = c(0.00719084498780675, 0.00989029529665065,
                 0.00576826003946358, 0.00781592130328325,
                 0.00298421642465751, 0.00166538046879564,
                 0.00048282648358605, 0.22975668514631000)
  
  # assign parameters
  hp <- parameters[1]
  ht <- parameters[2]
  hb <- parameters[3]
  hl <- parameters[4]
  hr <- parameters[5]
  he <- parameters[6]
  loss <- parameters[7]
  hd <- 0 # no diagonal transfer
  
  sz <- dim(on_wells)
  heat_rate <- parameters[8]
  
  # prepare generic convolution lookup table
  h_well_lut <- matrix(c(hd, hd, hl, hd, hd,
                         hd, hd, hl, hd, hd,
                         ht, ht, 0,  hb, hb,
                         hd, hd, hr, hd, hd,
                         hd, hd, hr, hd, hd), nrow=5)
  
  conv_internal <- matrix(c(hd, hp, hd,
                            hp, 0, hp,
                            hd, hp, hd), nrow=3)
  
  con_lut0 <- kronecker(matrix(1,8,12), conv_internal)
  
  # top + bottom
  top <- he*matrix(c(0,1,0),1,36)
  con_lut0[c(1, 24),] <- top
  
  # left + right
  left <- he*matrix(c(0,1,0),24,1)
  con_lut0[,c(1, 36)] <- left
  
  #
  con_lut_all = array(con_lut0,dim=c(24,36,sz[3]))
  
  for (i in 2:sz[3]) {
    con_lut <- con_lut0
    on_wells_c <- on_wells[,,i]
    h_wells <- which(on_wells_c > 0, arr.ind=TRUE)
    
    # Vectorized operations for heated wells
    rows <- h_wells[,1]
    cols <- h_wells[,2]
    indices <- cbind(rows, cols)
    
    if (!is_empty(indices)){
      for (w in 1:(length(indices)/2)){
        row = indices[w,1]
        col = indices[w,2]
        
        # check if heated well is at the edge
        if (row == 1 && col == 1){ 
          con_lut[2:4,2:4] = h_well_lut[3:5,3:5]
        }else if (row == 1 && col == 12){ #some problem with the else if loops
          con_lut[2:4,33:35] = h_well_lut[3:5,1:3]
        }else if (row == 8 && col == 1){
          con_lut[21:23,2:4] = h_well_lut[1:3,3:5]
        }else if (row == 8 && col ==12){
          con_lut[21:23,33:35] = h_well_lut[1:3,1:3]
        }else if (row == 1){
          col_min = col*3 - 3
          col_max = col*3 + 1
          con_lut[2:4,col_min:col_max] = h_well_lut[3:5,1:5]
        }else if (row == 8){
          col_min = col*3 - 3
          col_max = col*3 + 1
          con_lut[21:23,col_min:col_max] = h_well_lut[1:3,1:5]
        }else if (col == 1){
          row_min = row*3 - 3
          row_max = row*3 + 1 
          con_lut[row_min:row_max,2:4] = h_well_lut[1:5,3:5]
        }else if (col == 12){
          row_min = row*3 - 3
          row_max = row*3 + 1 
          con_lut[row_min:row_max,33:35] = h_well_lut[1:5,1:3]
        }else { #assign whole h_well_lut
          col_min = col*3 - 3
          col_max = col*3 + 1
          row_min = row*3 - 3
          row_max = row*3 + 1 
          con_lut[row_min:row_max,col_min:col_max] = h_well_lut
        }
      }
    }

    #loop over all wells to assign middle values
    for (r in c(1:8)){
      for (co in c(1:12)){
        conv = con_lut[(r*3-2):(r*3),(co*3-2):(co*3)]
        middle = -1*Reduce('+', conv)
        conv[2,2] = middle
        con_lut[(r*3-2):(r*3),(co*3-2):(co*3)] = conv 
      }
    }
    con_lut_all[,,i] = con_lut
  }
  
  # upsampling onwells and lookup table
  total_time = ceiling((sz[3]-1)*timestep)
  on_wells_1s = array(0, dim=c(sz[1],sz[2],total_time))
  con_lut_1s = array(0, dim=c(24,36,total_time))
  
  for (i in 1:(sz[3]-1)){
    step_range = (floor((i-1)*timestep+0.5)+1):(floor(i*timestep+0.5))
    on_wells_1s[,,step_range] = on_wells[,,i+1]
    con_lut_1s[,,step_range] = con_lut_all[,,i+1]
  }
  
  # preallocate
  timecourse_1s <- array(0, dim=c(8, 12, total_time))
  
  for (i in 2:total_time){
    pad =  padarray(timecourse_1s[,,i-1],c(1,1),0,'both')
    dtdt = zeros(8,12)
    con_lut_i = con_lut_1s[,,i]
    
    #loop over all the wells
    for (r in c(1:8)){
      for (co in c(1:12)){
        conv = con_lut_i[(r*3-2):(r*3),(co*3-2):(co*3)]
        neighbors = pad[(r):(r+2),(co):(co+2)]
        dtdt[r,co] = Reduce('+',conv*neighbors)-loss*(neighbors[2,2])
      }
    }
    
    timecourse_1s[,,i] = timecourse_1s[,,i-1]+dtdt
    
    on_wells_c <- on_wells_1s[,,i]
    h_wells <- which(on_wells_c > 0, arr.ind=TRUE)
    
    # Vectorized operations for heated wells
    rows <- h_wells[,1]
    cols <- h_wells[,2]
    indices <- cbind(rows, cols)
    
    if (!is_empty(indices)){
      #heat up heated wells
      for (hw in 1:(length(indices)/2)){
        cd = on_wells_c[indices[hw,1],indices[hw,2]] - timecourse_1s[indices[hw,1],indices[hw,2],i-1] #current difference from set-point
        if (cd > 0){
          timecourse_1s[indices[hw,1],indices[hw,2],i] = timecourse_1s[indices[hw,1],indices[hw,2],i]+heat_rate
        }
      }
    }

    # If we were passed a progress update function, call it and update progress
    if (is.function(updateProgress)) {
      text <- paste0("Model time: ", round(i/60,2), " min")

      updateProgress(value = i/sz[3], detail = text)
    }
  }
  
  # downsample 'timecourse_1s' to match initial desired output
  timecourse <- array(0, dim=sz)
  
  for (i in 1:(sz[3]-1)){
    step_sample = floor((i-1)*timestep+0.5)+1
    timecourse[,,i] = timecourse_1s[,,step_sample]
  }
  
  timecourse[,,sz[3]] = timecourse_1s[,,total_time]
  
  return(timecourse)
}
```
