## Supplementary material for "Multiplexed dynamic control of temperature to probe and observe mammalian cells": Files for running the thermoPlate GUI: SampleInputFile.R

```
SampleInputFile <- function(con){
  # To set temps use the amount of degrees above ambient temperature.
  OnTemp = array(c(
    c(10, 10, 10, 0, 0, 10, 10, 10), #1
    c(0, 0, 0, 0, 0, 0, 0, 0), #2
    c(0, 0, 0, 0, 0, 3, 0, 0), #3
    c(0, 0, 7, 0, 0, 0, 0, 0), #4
    c(0, 0, 0, 0, 0, 0, 0, 0), #5
    c(0, 0, 0, 0, 0, 0, 0, 0), #6
    c(0, 0, 0, 0, 0, 0, 0, 0), #7
    c(0, 0, 0, 1, 0, 0, 0, 0), #8
    c(0, 0, 0, 0, 0, 0, 0, 6), #9
    c(0, 8, 0, 0, 0, 0, 0, 0), #10
    c(0, 0, 0, 0, 0, 0, 0, 0), #11
    c(0, 0, 0, 0, 0, 0, 4, 0)  #12
    # A  B  C  D  E  F  G  H
  ), dim= c(8,12))
  
  OffTemp = array(c(
    c(2, 2, 2, 0, 0, 2, 2, 2), #1
    c(0, 0, 0, 0, 0, 0, 0, 0), #2
    c(0, 0, 0, 0, 0, 0, 0, 0), #3
    c(0, 0, 3, 0, 0, 0, 0, 0), #4
    c(0, 0, 0, 0, 0, 0, 0, 0), #5
    c(0, 0, 0, 0, 0, 0, 0, 0), #6
    c(0, 0, 0, 0, 0, 0, 0, 0), #7
    c(0, 0, 0, 9, 0, 0, 0, 0), #8
    c(0, 0, 0, 0, 0, 0, 0, 6), #9
    c(0, 4, 0, 0, 0, 0, 0, 0), #10
    c(0, 0, 0, 0, 0, 0, 0, 0), #11
    c(0, 0, 0, 0, 0, 0, 5, 0)  #12
    # A  B  C  D  E  F  G  H
  ), dim= c(8,12))
  
  OnTime = array(c(
    c(10, 10, 10, 0, 0, 10, 10, 10), #1
    c(0, 0, 0, 0, 0, 0, 0, 0), #2
    c(0, 0, 0, 0, 0, 4, 0, 0), #3
    c(0, 0, 5, 0, 0, 0, 0, 0), #4
    c(0, 0, 0, 0, 0, 0, 0, 0), #5
    c(0, 0, 0, 0, 0, 0, 0, 0), #6
    c(0, 0, 0, 0, 0, 0, 0, 0), #7
    c(0, 0, 0, 6, 0, 0, 0, 0), #8
    c(0, 0, 0, 0, 0, 0, 0, 6), #9
    c(0, 8, 0, 0, 0, 0, 0, 0), #10
    c(0, 0, 0, 0, 0, 0, 0, 0), #11
    c(0, 0, 0, 0, 0, 0, 4, 0)  #12
    # A  B  C  D  E  F  G  H
  ), dim= c(8,12))
  
  OffTime = array(c(
    c(10, 10, 10, 0, 0, 10, 10, 10), #1
    c(0, 0, 0, 0, 0, 0, 0, 0), #2
    c(0, 0, 0, 0, 0, 6, 0, 0), #3
    c(0, 0, 7, 0, 0, 0, 0, 0), #4
    c(0, 0, 0, 0, 0, 0, 0, 0), #5
    c(0, 0, 0, 0, 0, 0, 0, 0), #6
    c(0, 0, 0, 0, 0, 0, 0, 0), #7
    c(0, 0, 0, 8, 0, 0, 0, 0), #8
    c(0, 0, 0, 0, 0, 0, 0, 6), #9
    c(0, 8, 0, 0, 0, 0, 0, 0), #10
    c(0, 0, 0, 0, 0, 0, 0, 0), #11
    c(0, 0, 0, 0, 0, 0, 4, 0)  #12
    # A  B  C  D  E  F  G  H
  ), dim= c(8,12))
  
  # Create a new workbook
  wb <- createWorkbook()
  
  # Add sheets to the workbook for each array
  addWorksheet(wb, "OnTemp")
  writeData(wb, sheet = "OnTemp", x = OnTemp, rowNames = FALSE, colNames = FALSE)
  
  addWorksheet(wb, "OffTemp")
  writeData(wb, sheet = "OffTemp", x = OffTemp, rowNames = FALSE, colNames = FALSE)
  
  addWorksheet(wb, "OnTime")
  writeData(wb, sheet = "OnTime", x = OnTime, rowNames = FALSE, colNames = FALSE)
  
  addWorksheet(wb, "OffTime")
  writeData(wb, sheet = "OffTime", x = OffTime, rowNames = FALSE, colNames = FALSE)
  
  # Save the workbook to a file
  saveWorkbook(wb, con, overwrite = TRUE)
}
```
