## Supplementary material for "Multiplexed dynamic control of temperature to probe and observe mammalian cells": Files for running the thermoPlate GUI: wellPlateAnim.R

```
wellPlateAnim = function(heatModel_){
  # Generates the GIF of the simulation
  #covert errors to stroke thicknesses for plotting
  heatModel_ = heatModel_ %>% 
    mutate(errStroke = case_when(
      error == 0 ~ .5,
      error == 1 ~ 4
    ))
  
  p = ggplot(heatModel_) +
    geom_point(aes(x = ColNum, y = RowNum, fill = temp, color = error, stroke = errStroke),  size = 17, shape = 21, show.legend = FALSE)+
    scale_fill_gradient(low = "white", high = "purple", na.value = "white") +
    scale_color_steps(low = "black", high = "red")+
    scale_x_continuous(breaks = 1:12, limits = c(0.75, 12.25)) +
    scale_y_reverse(limits = c(8.25,0.75), breaks = 1:8, labels = LETTERS[1:8])+
    theme_bw() +
    theme(panel.grid.major = element_blank(),
          panel.grid.minor = element_blank(),
          axis.title.x=element_blank(),
          axis.title.y=element_blank(),
          axis.text.x = element_text(size = 12),
          axis.text.y = element_text(size = 12))+
    geom_text(aes(x = ColNum, y = RowNum, label = round(temp, digits =1), size = 0.2), show.legend = FALSE)+
    transition_time(time)+
    ggtitle("Time: {round(frame_time/60, 2)} min") #convert to minutes 
  
  anim = animate(p, renderer = gifski_renderer(), height = 400, width = 600, fps = 10)
  
  return(anim)
}
```
