## Supplementary material for "Multiplexed dynamic control of temperature to probe and observe mammalian cells": Files for running the thermoPlate GUI: wellPlatePlot.R

```
wellPlatePlot = function(wells_inputs, time_,ExperimentLength){
  # Makes the planned heat profile preview's input
  
  #change the 0s to NAs for plotting purposes. 
  dat = wells_inputs %>%   
    mutate(temp = 
             case_when(
               (temp == 0) & (start == 0) & (stop == ExperimentLength) ~ NA_real_,
               TRUE ~ temp
    ))
  
  #plot wellPlate over time with input data
  p = ggplot(wells_inputs) +
    geom_point(aes(x = ColNum, y = RowNum), size = 17, shape = 21,colour = "black", na.rm = TRUE)+
    geom_point(data = dat %>% filter (start <= time_ & time_ < stop), aes(x = ColNum, y = RowNum, fill = temp), size = 17, shape = 21, show.legend = FALSE, na.rm = TRUE)+
    scale_x_continuous(breaks = 1:12, limits = c(0.75, 12.25)) +
    scale_y_reverse(limits = c(8.25,0.75), breaks = 1:8, labels = LETTERS[1:8])+
    scale_fill_gradient(low = "white", high = "purple", limits = c(0,15), na.value="white") +
    theme_bw() +
    theme(panel.grid.major = element_blank(),
          panel.grid.minor = element_blank(),
          axis.title.x=element_blank(),
          axis.title.y=element_blank(),
          axis.text.x = element_text(size = 12),
          axis.text.y = element_text(size = 12))+
    geom_text(data = dat %>% filter (start <= time_ & time_ < stop), aes(x = ColNum, y = RowNum, label = temp, size = 0.2), show.legend = FALSE, na.rm = TRUE)+
    #transition_time(time)+
    ggtitle(paste("Time:", time_, "min", sep = " "))
  
  return(p)
}
```
