## Supplementary figures and images for "Multiplexed dynamic control of temperature to probe and observe mammalian cells"

### simPlaceHolder.png

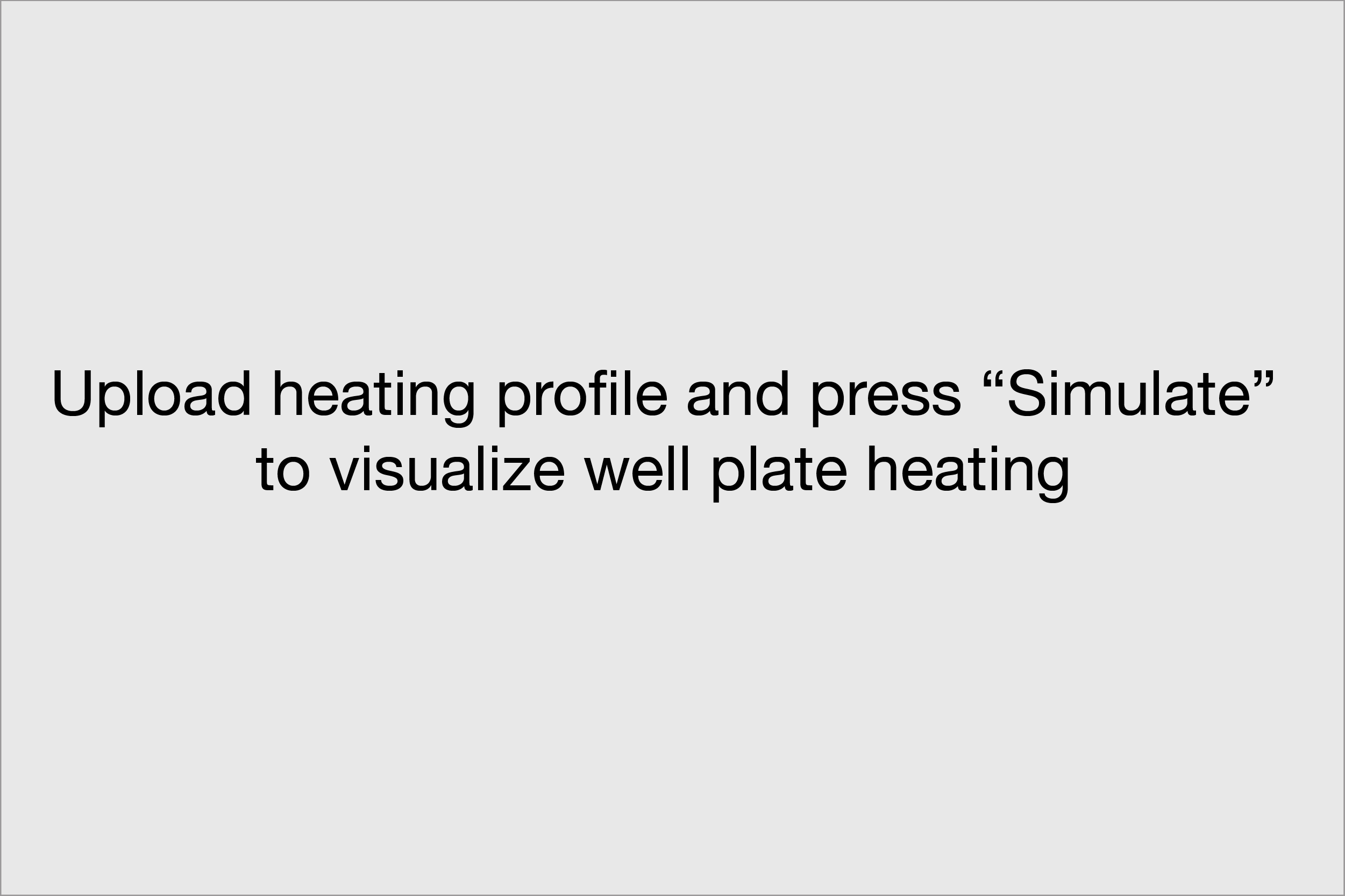

### test.gif

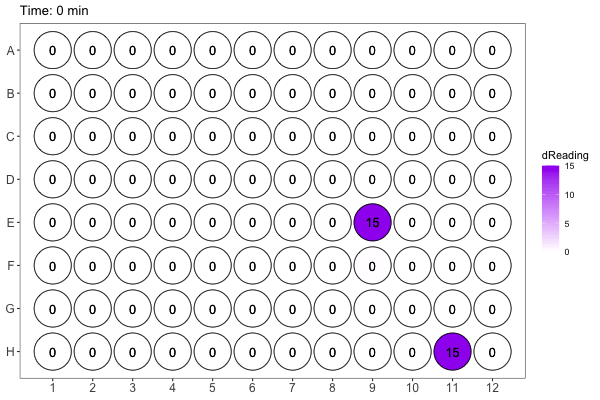
